# MPDB: a unified multi-domain protein structure database integrating structural analogue detection

**DOI:** 10.1101/2021.10.27.466092

**Authors:** Chun-Xiang Peng, Xiao-Gen Zhou, Yu-Hao Xia, Yang Zhang, Gui-Jun Zhang

## Abstract

With the development of protein structure prediction methods and biological experimental determination techniques, the structure of single-domain proteins can be relatively easier to be modeled or experimentally solved. However, more than 80% of eukaryotic proteins and 67% of prokaryotic proteins contain multiple domains. Constructing a unified multi-domain protein structure database will promote the research of multi-domain proteins, especially in the modeling of multi-domain protein structures. In this work, we develop a unified multi-domain protein structure database (MPDB). Based on MPDB, we also develop a server with two functional modules: (1) the culling module, which filters the whole MPDB according to input criteria; (2) the detection module, which identifies structural analogues of the full-chain according to the structural similarity between input domain models and the protein in MPDB. The module can discover the potential analogue structures, which will contribute to high-quality multi-domain protein structure modeling.

## INTRODUCTION

Multi-domain proteins are an important class of proteins, and consist of more than one unique folding units that are connected into a single chain. Many biological functions rely on the interaction of different domains [1]. For example, the ribose-binding protein in bacterial transport and chemotaxis, the ligand-binding site is located in a pocket formed by two domains. However, only one third of structures in the Protein Data Bank (PDB) contain multiple domains. There is an ever-growing interest to research multi-domain proteins when more and more single-domain proteins are experimentally solved or modeled by high-accuracy deep-learning protein structure prediction algorithms such as AlphaFold2 [2], RoseTTAFold [3], D-I-TASSER [4] etc.

There are some databases for protein domains. The CATH (Class, Architecture, Topology, Homology) is a database which classifies the protein domains into families based on structures deposited in the Protein Data Bank (PDB) [5]. SCOP (Structural Classification of Proteins) database is a manual classification of protein structural domains according to the structural and evolutionary relationships [6], and SCOPe is an extended database of SCOP, which classifies protein domains through a combination of automation and manual curation based on the similarity of structures and amino acid sequences. Both of them are widely used in analyzing protein sequence, structure, function, and evolution and in developing various bioinformatics tools. ECOD (Evolutionary Classification of protein Domains) groups domains primarily by evolutionary relationships (homology) [7], which helps homology-based structure and function prediction and protein annotation by providing a pre-compiled search database. These databases are protein structural domain databases, which have classified protein individual domains according to different information and definitions. However, to the best of our knowledge, there is no reliable and comprehensive structure database in the literature for the full-chain multi-domain proteins, despite the increasing interest of multi-domain structure and function modeling. In addition, the quality of protein structure predictions is also assessed mainly on the individual domains in community-wide CASP experiments [8]. Similarly, most computational approaches are only optimized for single-domain structure predictions [9]. These reasons may lead to the research of protein structure focusing more on domain or single-domain protein, especially in protein structure prediction. Nevertheless, there is an indisputable fact that most proteins consist of multiple domains and the research of full-chain structure of multi-domain proteins is thus a crucial step in elucidating their functions and designing new drugs to regulate these functions.

Templates are important for modeling protein structure. Since the conformational information from template is much more reliable than that from elsewhere (especially when the target protein and the template are highly homologous), the prediction accuracy of template-based method is generally higher than other methods, which makes it highly popular in practical applications [10]. Currently, templates can be identified from PDB by profile-profile alignment or sequence-structure threading method, such as HHblits [11], HHsearch [12], LOMETS2 [13], DeepThreader [14] and so on. However, some proteins are formed by duplication, divergence and recombination of domains [15]. Meanwhile, the structure of single-domain protein can be relatively easier to be experimentally solved or computationally modelled, especially with the revolutionary success of AlphaFold2 for single-domain protein structure modeling [2]. Therefore, inferring structural analogues of the full-chain based on the domain structures may be more critical for protein full-chain modeling in some cases, especially for those proteins with only partial domains known.

In this work, our main objective is to develop a unified information portal for the researchers interested in multi-domain proteins. We have attempted to collect as many multi-domain proteins as possible from PDB. Here, a unified multi-domain protein structure database (MPDB) is developed based on the PDB library by using the DomainParser and the domain knowledge defined in the CATH and SCOPe database. In addition, we further develop two functional modules based on MPDB. The culling module filters the whole MPDB according to input criteria, and the structural analogues detection module identifies structural analogues of the full-chain from MPDB through structural alignment. The online server for MPDB is freely available at http://zhanglab-bioinf.com/MPDB.

## RESULTS

Until October, 2021, MPDB contains 48,224 multi-domain proteins, in which 37,495 proteins have 2 domains, 7,538 proteins have 3 domains, 2,182 proteins have 4 domains, 1,009 proteins have more than 4 domains. The MPDB will be updated periodically following the PDB library. The statistics of the number of domains in the MPDB are shown in Figure 1 (a).

**Figure 1.**
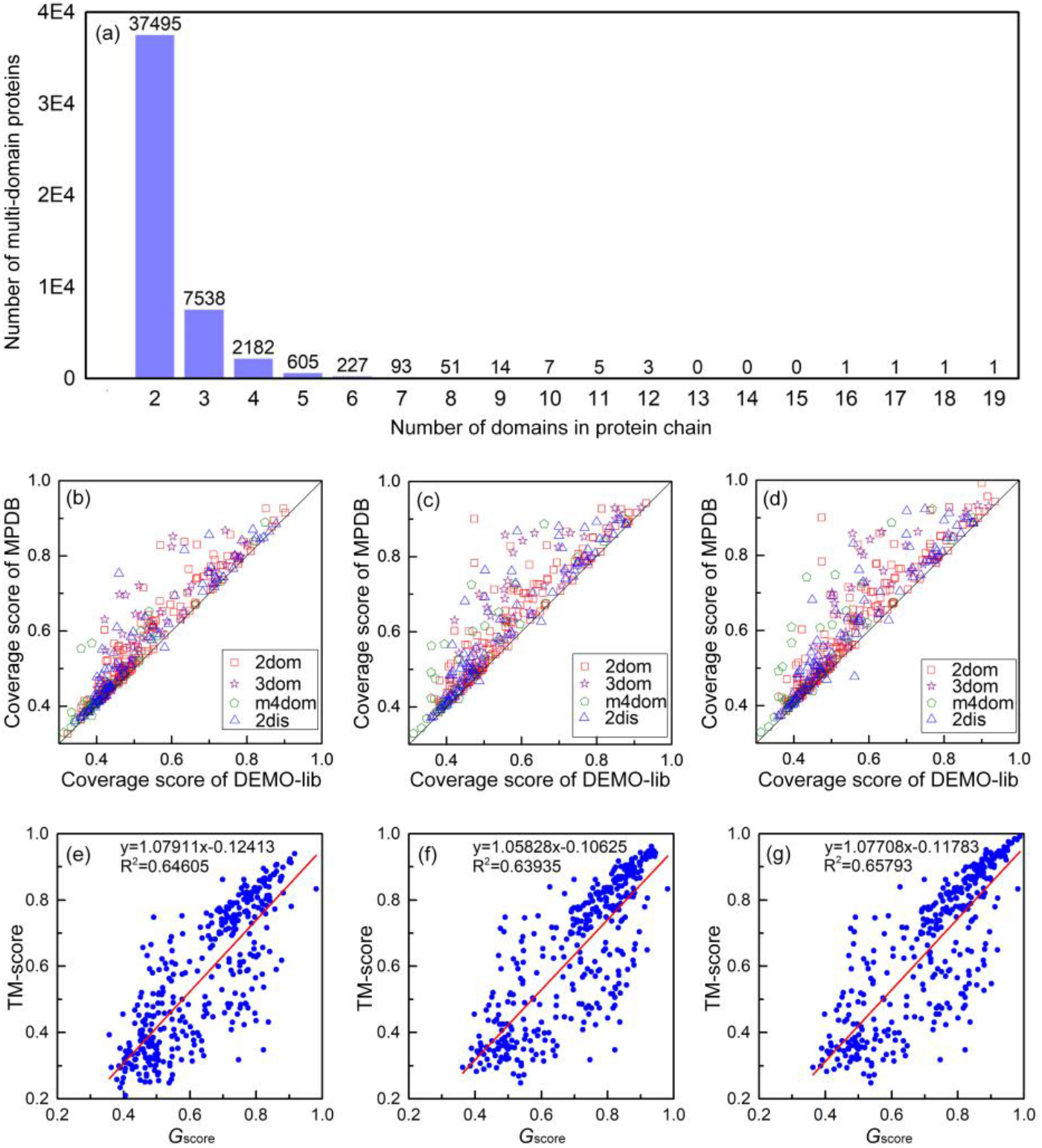
(a) Distribution of different multi-domain proteins in MPDB. (b), (c) and (d) represent head-to-head comparison between the coverage score of the test proteins in DEMO-lib and MPDB under the condition that the sequence identity between the test protein and the proteins in MPDB and DEMO-lib is less than 30%, 50% and 70%, respectively. The x-axis represents the coverage score of the test proteins on the DEMO-lib, where DEMO-lib represents the template library used in DEMO. The y-axis represents the coverage score of the test proteins on the MPDB. (e), (f) and (g) represent the distribution of *G*_score_ and TM-score under sequence identity less than 30%, 50% and 70%, respectively. For each blue dot, the x-axis represents *G*_score_ of the structural analogue with the highest *G*_score_ in the corresponding test protein, and the y-axis represents the TM-score between the native structure of the test protein and the detected structural analogue. The least-squares linear fit and R-square are listed in (e), (f) and (g).

### Coverage of multi-domain proteins in the MPDB

In order to test the coverage of data in the MPDB, we compare the coverage of MPDB and that of template libraries used in our previous developed DEMO [9] for multi-domain protein structures assembly. Here, template library used in DEMO is denoted as DEMO-lib, and we use all of the 356 proteins from DEMO benchmark dataset as the test targets. The details of the 356 test proteins are shown in Table S1, including PDB ID, protein length and the number of domains. In the MPDB and DEMO-lib, TM-align [16] is employed to calculate the TM-score [17] between each test protein and the proteins in the two databases, respectively. For each test protein, the average TM-score of top 10 templates with the highest TM-score is considered as the coverage score of the test protein in the database.

Under the sequence identity cutoff of 30%, the average coverage score of MPDB is 0.57, which is 7.5% higher than that of DEMO-lib (0.53) for the 166 2-domain (2dom) proteins. The average coverage score of MPDB is 0.56, which is 12.0% higher than that of DEMO-lib (0.50) for the 69 3-domain (3dom) proteins. For the 40 ≥ 4-domain (m4dom) proteins, the average coverage score of MPDB is 0.48, which is 6.7% higher than that of DEMO-lib (0.45). The average coverage score of MPDB is 0.56 for the 81 discontinuous-domain (2dis) proteins, which is 5.7% higher than that of DEMO-lib (0.53). The *P*-value of the coverage scores of MPDB and that of DEMO-lib is 2.04E-04, indicating that there is a significantly difference between the coverage score of MPDB and DEMO-lib. Under the sequence identity cutoff of 50%, the average coverage score of MPDB is 0.60, which is 9.1% higher than that of DEMO-lib (0.55) for the 356 test proteins. When using the sequence identity cutoff of 70%, the average coverage score of MPDB is 0.61, which is 10.9% higher than that of DEMO-lib (0.55). The detailed coverage score for each test protein and the summaries of different databases are listed in Supplementary Tables S2-S7. The head-to-head comparisons between the coverage score of each test protein in DEMO-lib and MPDB are shown in Figures. 1 (b), (c) and (d), which show that the MPDB covers significantly more multi-domain protein structures than DEMO-lib and the significantly difference between the protein coverage of MPDB and DEMO-lib.

### Performance of structural analogue detection and application

There are some proteins that the sequence identity between them is low, but they have similar topologies. These proteins may be more important for structural modeling of multi-domain proteins, especially for those proteins with only partial domains determined. Therefore, we develop a structural analogue detection module, which is introduced in method section. Here, we tested the performance of the structural analogue detection as follow.

In the 356 test proteins, the structural analogue detection module is used to detect structural analogues of the full-chain according to the individual domain models, and the structural analogue with the highest *G*_score_ is used to calculate TM-score between the analogue and native structure of target protein. Here, *G*_score_ is the harmonic mean of the TM-scores between the domain models of the target protein and the multi-domain protein in MPDB. *G*_score_ has a value range of (0, 1], where 1 indicates a perfect match between domain models and the structural analogues of the full-chain, and the details of *G*_score_ are described in the method section. When the sequence identity is less than 30%, the average TM-score of the 356 structural analogues with the highest *G*_score_ is 0.56, where 192 cases have the highly similar global topologies to the structural analogues with TM-score > 0.5. Especially, the TM-score between structural analogues with the highest *G*_score_ and corresponding native structures of the proteins 3qtdA with 2 domains, 1zpuA with 3 domains, and 1z1wA with 4 domains are 0.94, 0.91 and 0.87, while the sequence identity between them is 29%, 26% and 24%, respectively. When the sequence identity is less than 50% and 70%, the average TM-score of the structural analogues is increased to 0.64 and 0.65, and the number of structural analogues with TM-score > 0.5 to the full-chain structures is 239 and 247, respectively. The detailed TM-score of structural analogue for each test protein in different sequence identity cutoff is shown in Supplementary Tables S8 and S9. Under the different sequence identity cutoff, the average TM-scores of structural analogues are always more than 0.5, indicating that the detected structural analogues can describe the global topological structure of the full-chain for the majority of multi-domain proteins in the test set.

In addition, in order to prove the rationality of *G*_score_, the Pearson correlation coefficient between *G*_score_ of the top 1 structural analogue detected and TM-score of the structural analogue to the native structure is calculated under the sequence identity cutoff of 30%, 50% and 70%, respectively. For the 356 test proteins, the *G*_score_ of the top 1 structural analogue is shown in Table S10. The distributions of *G*_score_ and TM-score are shown in Figures. 1 (e), (f) and (g). Under sequence identity cutoff of 30%, 50% and 70%, the Pearson correlation coefficients between the *G*_score_ and TM-score are 0.80, 0.80 and 0.81, respectively. The results show that the designed *G*_score_ can effectively evaluate the structural analogues of full-chain.

The structural analogues are crucial for the modeling of multi-domain protein structures and the discovery of potential analogues of domains, especially for those proteins with only partial domains determined. For example, DEMO [9] constructs multi-domain protein structures through docking-based domain assembly simulations based on the structural analogous templates. Here, we show the application of structural analogues with two examples. In Figure 2 (a), the individual domains of 4uwhA are treated as the query domains, and the structural analogue (5uk8A) with the highest *G*_score_ is detected from MPDB under the sequence identify cutoff of 30%. The structural analogue (5uk8A) is similar with the native structure of 4uwhA in structure topology, and the TM-score between 5uk8A and 4uwhA is 0.85. The analogue 5uk8A and experimental domains of 4uwhA are input into DEMO as the template and query domains, respectively, and a final model with TM-score = 0.99 to the native structure is generated, which reinforces the importance of an analogous template with high quality for the multi-domain protein structure assembly. In addition, structural analogues can be used to model unknown domain structure (or full-chain structure), when only partial domains of a protein are known. In this case, structural analogues may be better than homologous templates, because the protein structure is much more conservative than sequence. Figure 2 (b) shows the protein 3ob8A with 5 domains, where we assume the largest domain (312 amino acids) is unknown. Based on the 4 known domains, the structural analogue 1f4aA (sequence identity = 32%) is detected. The TM-score between the full-chain native structure of 3ob8A and 1f4aA is 0.89, and the native structure of 3ob8A and 1f4aA has a TM-score of 0.86 in the region of unknown domain of 3ob8A. This case indicates that structural analogues are important for the full-chain structure and domain modeling even when only partial region has known structure in some cases.

**Figure 2.**
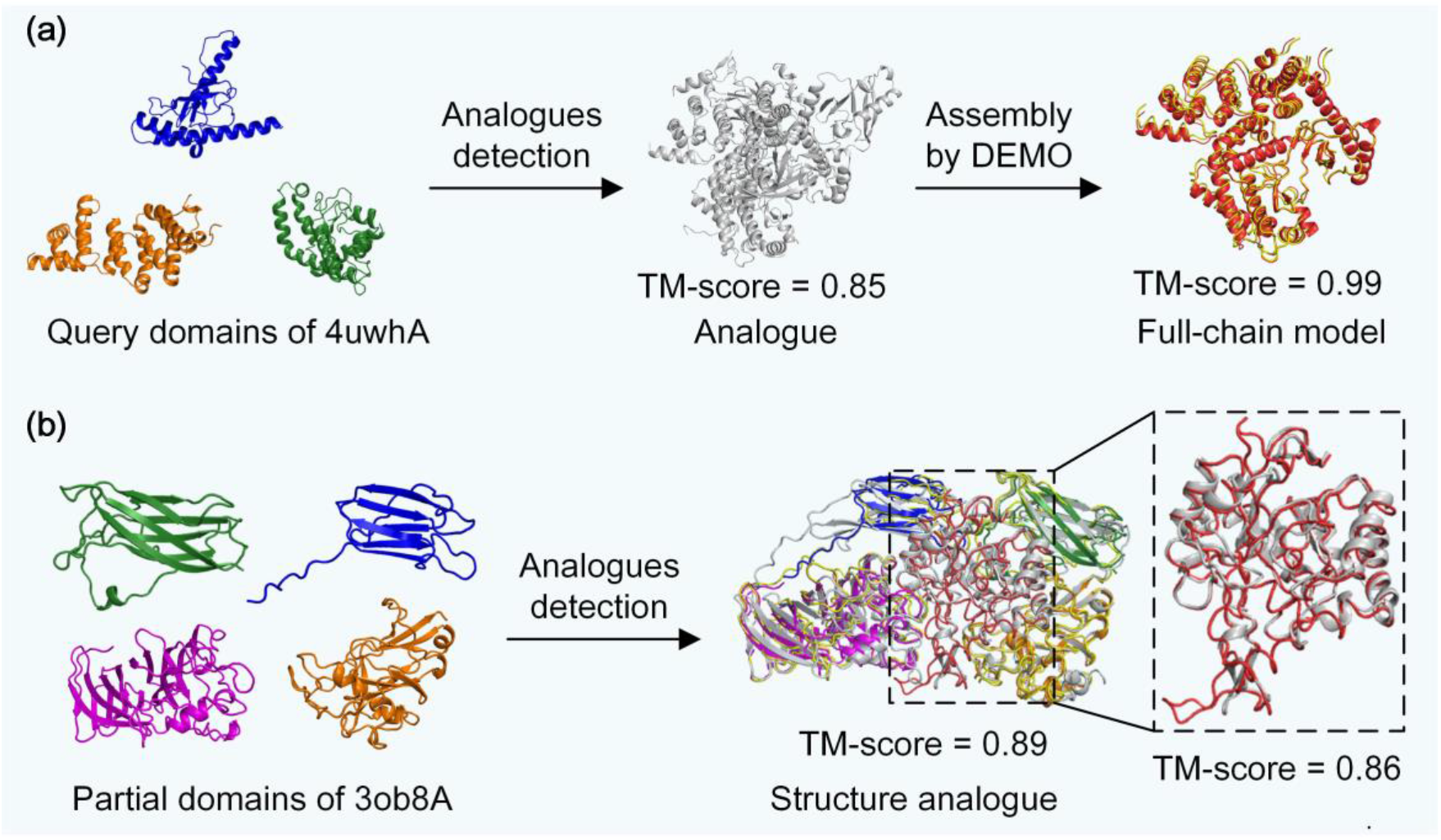
Application of the structural analogues. (a) Structural analogues for multi-domain protein structure assembly using individual domain models. The orange, blue and green models represent the input individual domain structures. The gray model is the detected structural analogue with the highest *G*score. The yellow and red models represent the native structure and final assembled full-chain structure, respectively. (b) The discovery of potential structural analogues of full-chain. The orange, blue, green and magenta models represent the input individual domain structures. The yellow and gray models represent the native structure of the target (3ob8A) and the structural analogue of full-chain, respectively. Red region represents the native structure of unknown domain.

### Online database

MPDB is built based on the Spring Boot (version 2.1.4) framework, Bootstrap 4 and Thymeleaf (version 3.0.11) template engine and runs on the tomcat server embedded in Spring Boot with MySQL (version 8.0.22) as its database engine. The NGL Viewer is used to display protein structures in three dimensions [18]. The architecture of MPDB web server is shown in Supplementary Fig. S1. MPDB can be used online without registration, and is optimized for task execution under high concurrent requests.

As shown in Supplementary Fig. S2, the top of the webserver of MPDB introduces the construction method of MPDB and the current number and distribution of multi-domain proteins in MPDB. In the middle of webserver, there are 2 function buttons provided to switch to different function modules. In the download part, we provide all structures in current MPDB and the domain boundary information. In the news part, we mainly record the updated information of MPDB.

In the structural analogue detection module, the mandatory inputs for the web server are individual domain structure files in PDB format and target sequence, where the sequence is used to calculate sequence identity. In the absence of an experimentally solved structure, the user can use models generated by the structure prediction tools, such as AlphaFold2 [2], D-I-TASSER [4], RoseTTAFold [3], CoDiFold [19] or CGLFold [20]. Upon job completion, the user will be notified by an email with two links to the results display page and results download page on the structural analogue detection module, respectively. The results display page shows an ordered list of the top-ten structural analogues with the highest *G*_score_ from the MPDB. The structural analogues are displayed in three dimensions by the NGL Viewer, which allows users to rotate the pictures, and there is a link is given to the URL addresses for downloading the analogue PDB structure. The structural analogue is shown together with the *G*_score_, TM-score of each input domain aligned on the analogue, aligned length between input sequence and structural analogue sequence, sequence identical length and sequence identity, where the aligned length, identical length and sequence identity are calculated by NW-align program (http://zhanglab.dcmb.med.umich.edu/NW-align). In the results download page, a compressed file containing structures of top 200 multi-domain protein structural analogues with the highest *G*_score_ and an analogues information file including the information mentioned above can be downloaded. In Supplementary Fig. S3 and Fig. S4, we show an example of the structural analogue detection in the MPDB server.

In the culling module, users need to input the criteria including protein chain length, resolution, number of domains, R-factor and sequence identity of multi-domain protein, and then the module retrieves whole MPDB according to the criteria. After the retrieval is finished, the module sends the user an email that includes a link to the resulting files that the user can then download the multi-domain proteins satisfying the criteria and the corresponding information file recording the protein name, chain length, experimental measurement methods, resolution, R-factor, number of domains, and methods of defining domain boundary and domain boundary. In Supplementary Fig. S5 and Fig. S6 we show an example of multi-domain proteins culling in the MPDB server.

## CONCLUSION

We develop a unified multi-domain protein structure database, termed MPDB. Based on MPDB, a multi-domain proteins culling module is developed, which filters the whole MPDB according to user’s input criteria including protein chain length, resolution, number of domains, R-factor and sequence identity of multi-domain protein, and then provides dataset of protein structures and related information that satisfy the criteria. Users can conveniently retrieve results for their protein(s) of interest. In addition, we develop the structural analogue detection module, which is crucial for the modeling of multi-domain protein structures and the discovery of potential structural analogues of the full-chain, especially for those proteins with only partial domains determined. For the 356 test proteins, the average TM-score of the structural analogues detected is 0.56, 0.64 and 0.65 and the number of structural analogues which have similar global topology (TM-score > 0.5) to the target is 192, 239 and 247, when the sequence identity cutoff is set to 30%, 50% and 70%, respectively. The results show that the detected structural analogues can describe the topological structure for the majority of multi-domain proteins in the test set. The structural analogues detection method may be extended to the detection of structural analogue of protein complex, which may be contributed to the protein complex modeling, and this is also the next direction of our group. We hope that a high-quality and up-to-date multi-domain structure database will have critical importance in facilitating investigations relevant to multi-domain proteins.

## METHODS

### Procedure for database construction

The pipeline of MPDB is shown in Figure. 3 (a). The collection and processing of multi-domain protein data in MPDB are as follows: (1) CD-HIT [21] is used to remove redundancy of protein structures with a sequence identity cutoff 100% in PDB, and then protein structures with sequence identity less than 100% are fetched from PDB; (2) DomainParser [22] is next used to determine whether these proteins are multi-domain proteins or not; (3) the single-domain proteins determined by DomainParser are further confirmed by CATH and SCOPe on whether they are multi-domain proteins. All the multi-domain proteins selected in the above 3 steps are finally collected to construct the MPDB.

**Figure. 3.**
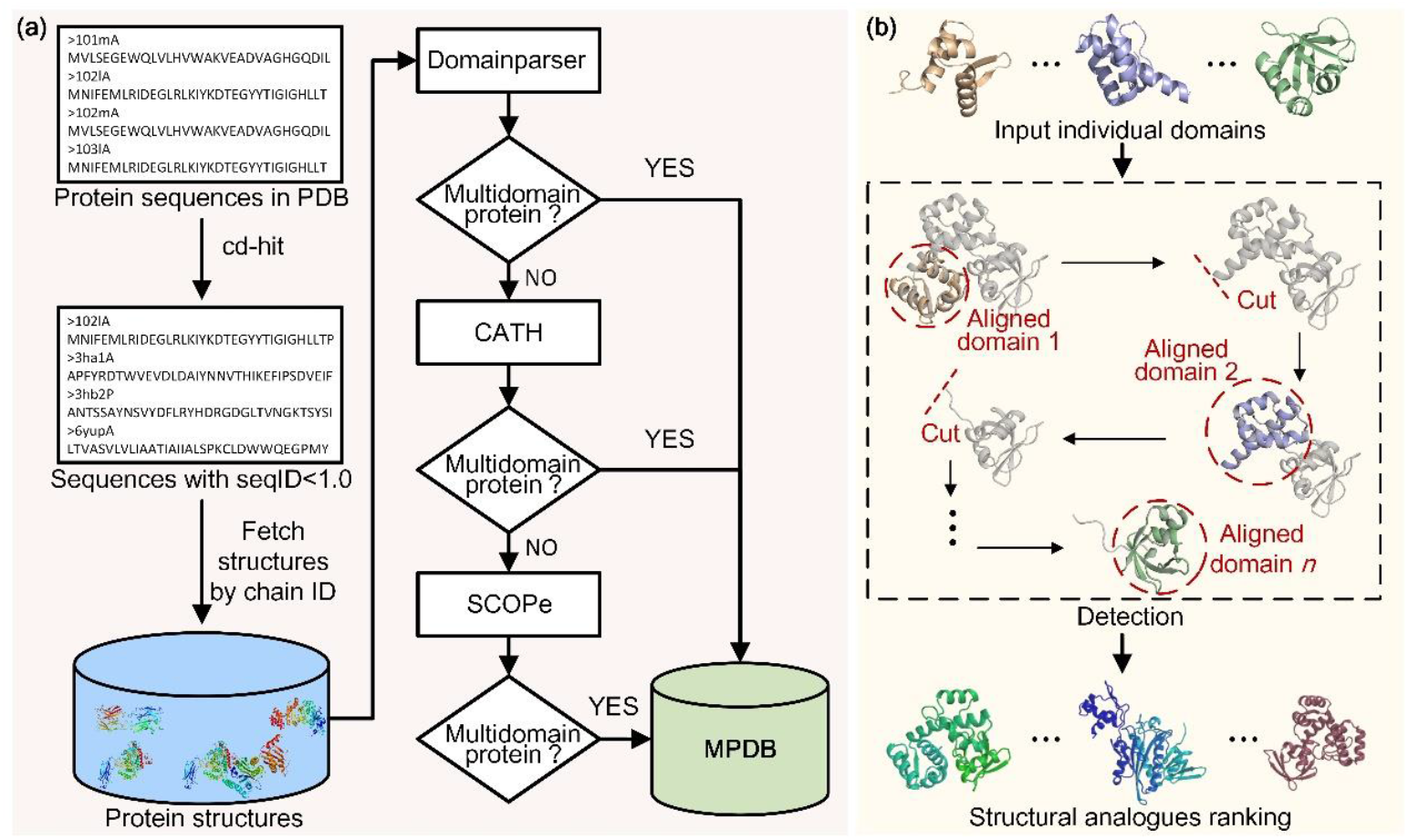
(a) Pipeline of MPDB construction method. (b) Illustration of structural analogues detection.

### Structural analogues detection

Based on MPDB, we developed a module to detect multi-domain protein structural analogues. The illustration of structural analogues detection is shown in Figure. 3 (b). The input individual domains are aligned on each protein in MPDB by TM-align [16, 17], with no overlap allowed in the alignments of different domains. This can prevent structurally similar domains from being matched to the same part of the protein, thereby improving the quality of structural analogue. A global score *G*_score_ is designed to evaluate the structural similarity between domain models and the protein in MPDB. The top 200 multi-domain protein structural analogues with the highest *G*_score_ are output. *G*_score_ is defined as follows:

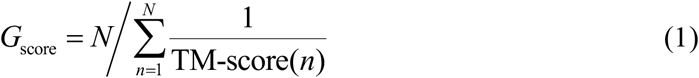

where TM-score(*n*) is the TM-score between the *n*-th domain of the target and the template protein in MPDB after aligning the domain on the template protein by TM-align. *N* is the number of query individual domains.

### Data retrieval

For MPDB, it is often the case that additional criteria are desirable, such as resolution, length, sequence identity or number of domains cutoffs. Inspired by PISCES [23], we developed a module for retrieving multi-domain proteins, which filters the whole MPDB according to user’s input criteria including protein chain length, resolution, number of domains, R-factor and sequence identity of multi-domain protein, and then provides the protein structures and related information that satisfy the criteria.

These criteria are applied first the list of MPDB entries. The result is a list of MPDB chains that fit the criteria that can then be culled according to mutual sequence identity. PSI-BLAST is used to calculate the sequence identity, and the details can be found in Supplementary text S1.

## Declaration of competing interest

The authors declare that they have no known competing financial interests or personal relationships that could have appeared to influence the work reported in this paper.

## Acknowledgments

We thank Zhongze Yu and Fenqi Ge for helpful discussions. This work has been supported in part by the National Nature Science Foundation of China (No. 62173304 and No. 61773346), the Key Project of Zhejiang Provincial Natural Science Foundation of China (No. LZ20F030002), the National Key Research and Development Program of China (No. 2019YFE0126100), the National Institute of General Medical Sciences (GM136422, S10OD026825), the National Institute of Allergy and Infectious Diseases (AI134678), and the National Science Foundation (IIS1901191, DBI2030790, MTM2025426).

## Author’s Contributions

G.Z. and Y.Z. designed the research, C.P., X.Z. and Y.X. performed the research, C.P., X.Z. and G.Z. wrote the manuscript, and all authors read and approved the final manuscript.

## Notes

### Competing Interest Statement

The authors have declared no competing interest.

## References

[1] Zhou, X., et al. (2020). Progressive and accurate assembly of multi-domain protein structures from cryo-EM density maps. bioRxiv, 2020.2010.2015.340455.

[2] Jumper, J., et al. (2021). Highly accurate protein structure prediction with AlphaFold. Nature, 596, 583–589.

[3] Baek, M., et al. (2021). Accurate prediction of protein structures and interactions using a three-track neural network. Science, 373, 871–876.

[4] Zheng, W., et al. (2021). Protein structure prediction using deep learning distance and hydrogen-bonding restraints in CASP14. Proteins, DOI:10.1002/prot.26193.

[5] Lam, S. D., et al. (2016). Gene3D: expanding the utility of domain assignments. Nucleic Acids Res., 44, D404–D409.

[6] Chandonia, J.-M., Fox, N. K. & Brenner, S. E. (2017). SCOPe: Manual Curation and Artifact Removal in the Structural Classification of Proteins – extended Database. J. Mol. Biol., 429, 348–355.

[7] Cheng, H., et al. (2014). ECOD: An Evolutionary Classification of Protein Domains. PLoS Comput. Biol., 10, e1003926–e1003926.

[8] Kinch, L. N., Kryshtafovych, A., Monastyrskyy, B. & Grishin, N. V. (2019). CASP13 target classification into tertiary structure prediction categories. Proteins, 87, 1021–1036.

[9] Zhou, X., Hu, J., Zhang, C., Zhang, G. & Zhang, Y. (2019). Assembling multidomain protein structures through analogous global structural alignments. Proc. Natl. Acad. Sci. USA, 116, 15930.

[10] Deng, H., Jia, Y. & Zhang, Y. (2017). Protein structure prediction. Int J Mod Phys B, 32, 1840009.

[11] Remmert, M., Biegert, A., Hauser, A. & Söding, J. (2012). HHblits: Lightning-fast iterative protein sequence searching by HMM-HMM alignment. Nature Methods, 9, 173–175.

[12] Söding, J. (2004). Protein homology detection by HMM–HMM comparison. Bioinformatics, 21, 951–960.

[13] Zheng, W., et al. (2019). LOMETS2: improved meta-threading server for fold-recognition and structure-based function annotation for distant-homology proteins. Nucleic Acids Res., 47, W429–W436.

[14] Zhu, J., Wang, S., Bu, D. & Xu, J. (2018). Protein threading using residue co-variation and deep learning. Bioinformatics, 34, i263–i273.

[15] Vogel, C., Bashton, M., Kerrison, N. D., Chothia, C. & Teichmann, S. A. (2004). Structure, function and evolution of multidomain proteins. Curr. Opin. Struct. Biol., 14, 208–216.

[16] Zhang, Y. & Skolnick, J. (2005). TM-align: a protein structure alignment algorithm based on the TM-score. Nucleic Acids Res., 33, 2302–2309.

[17] Zhang, Y. & Skolnick, J. (2004). Scoring function for automated assessment of protein structure template quality. Proteins, 57, 702–710.

[18] Rose, A. S. & Hildebrand, P. W. (2015). NGL Viewer: a web application for molecular visualization. Nucleic Acids Res., 43, W576–W579.

[19] Peng, C., Zhou, X. & Zhang, G. (2020). De novo Protein Structure Prediction by Coupling Contact with Distance Profile. IEEE/ACM Trans. Comput. Biol. Bioinf., DOI:10.1109/TCBB.2020.3000758.

[20] Liu, J., Zhou, X.-G., Zhang, Y. & Zhang, G.-J. (2020). CGLFold: a contact-assisted de novo protein structure prediction using global exploration and loop perturbation sampling algorithm. Bioinformatics, 36, 2443–2450.

[21] Fu, L., Niu, B., Zhu, Z., Wu, S. & Li, W. (2012). CD-HIT: Accelerated for clustering the next-generation sequencing data. Bioinformatics, 28, 3150–3152.

[22] Xu, Y., Xu, D. & Gabow, H. N. (2000). Protein domain decomposition using a graph-theoretic approach. Bioinformatics, 16, 1091–1104.

[23] Wang, G. & Dunbrack, R. L. (2003). PISCES: a protein sequence culling server. Bioinformatics, 19, 1589–1591.

