## Supplementary Information for "MPDB: a unified multi-domain protein structure database integrating structural analogue detection"

**Supplementary Figures**

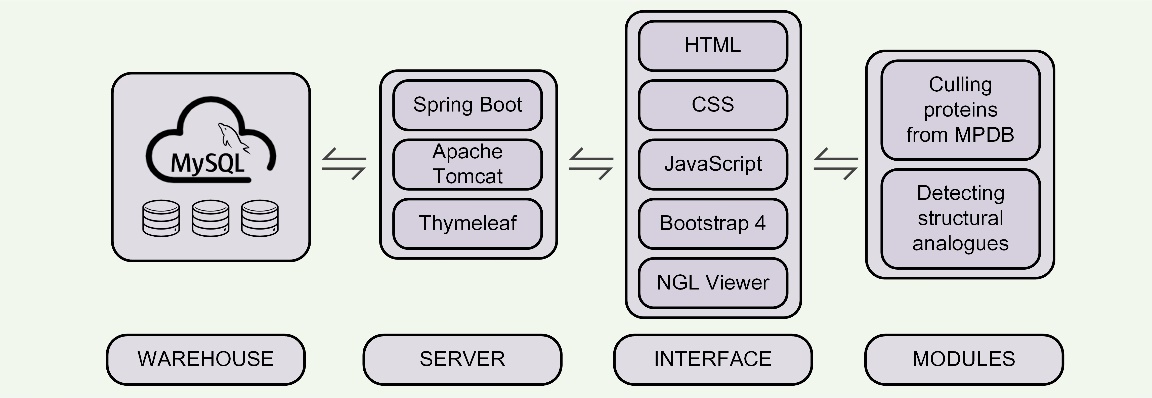

**Fig. S1.** The architecture of MPDB web server.

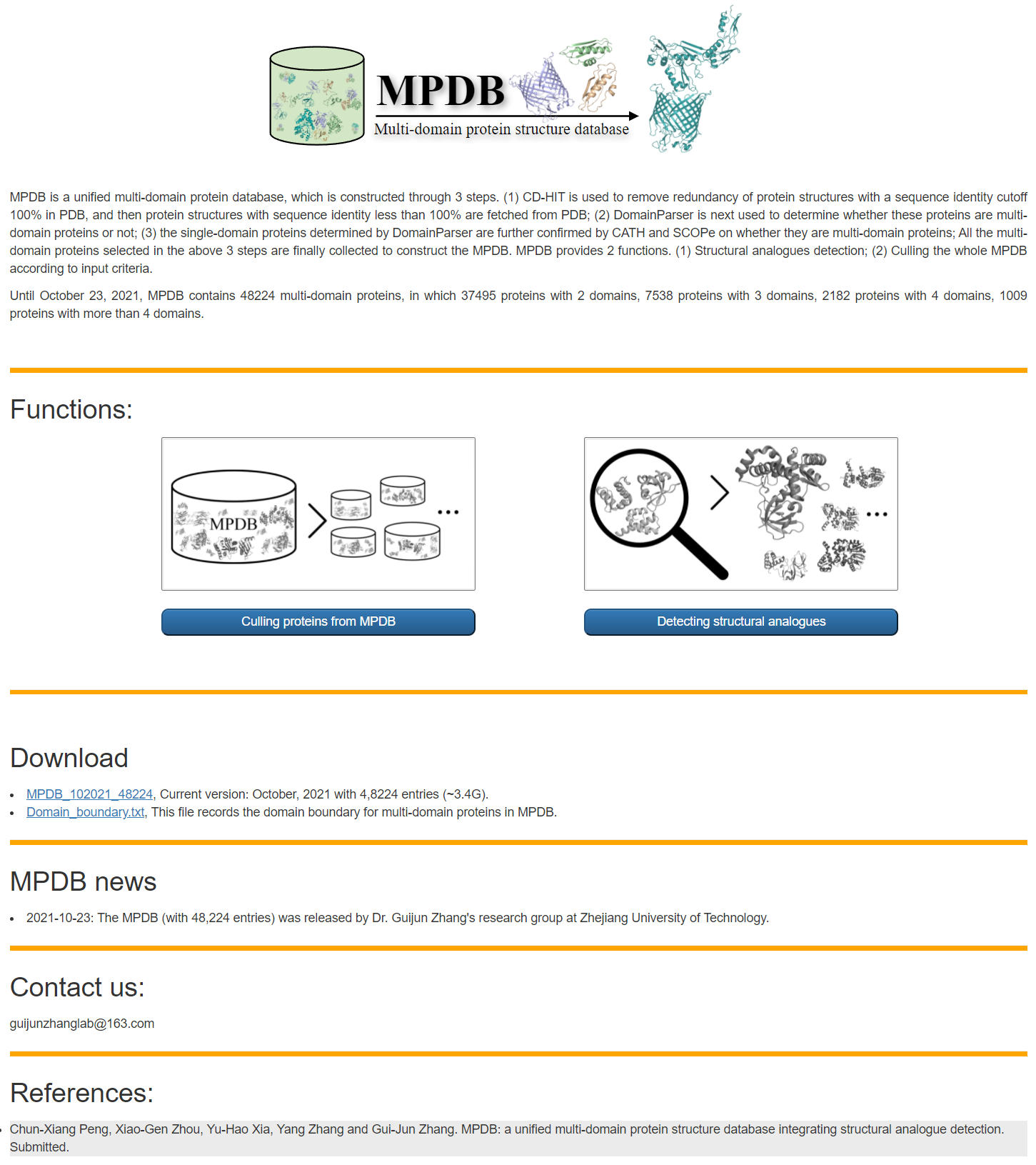

**Fig. S2.** The homepage of MPDB web server.

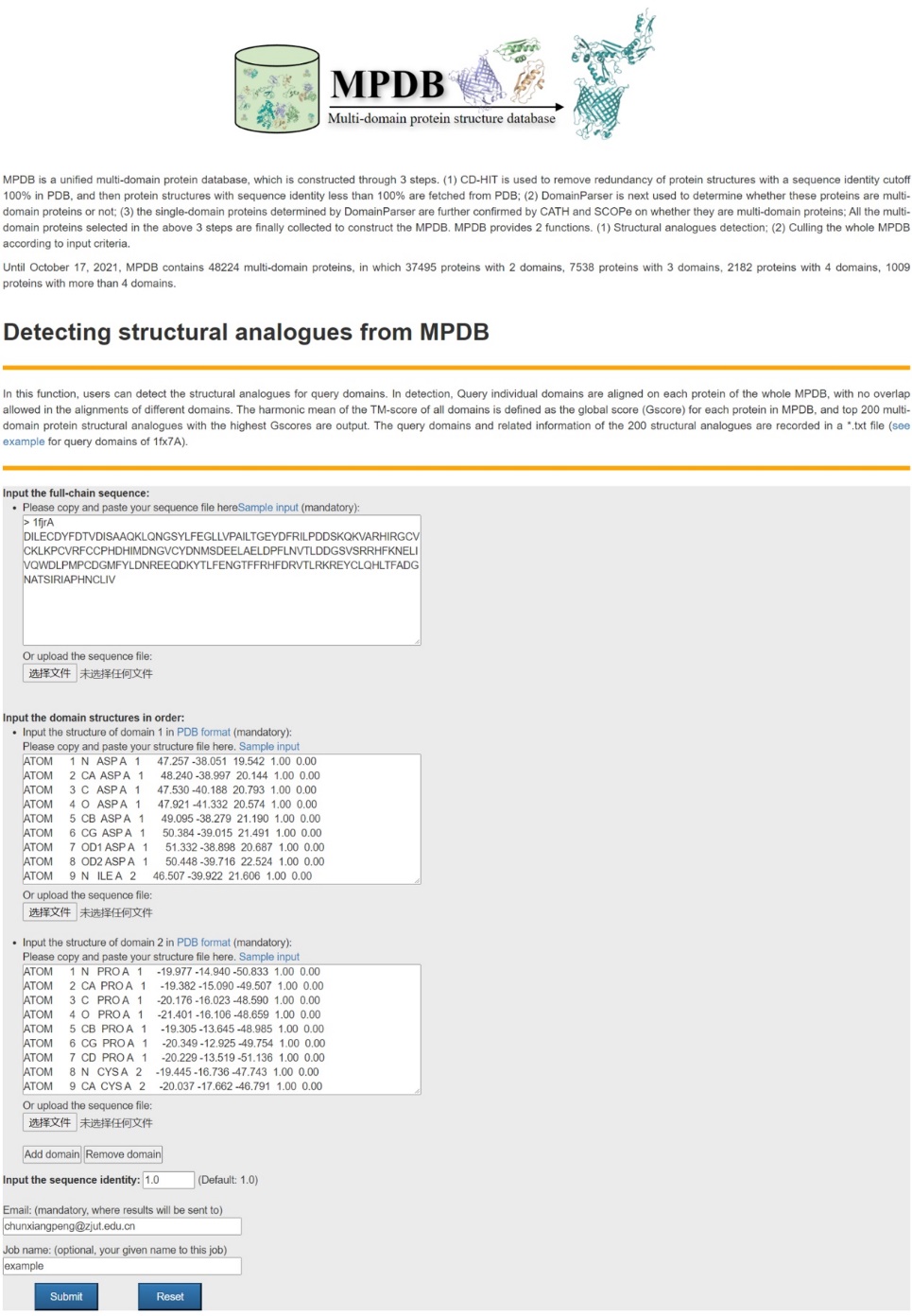

**Fig. S3.** The input page of structural analogue detection module in the MPDB web server. According to the target sequence, query domain models and sequence identity input by the users, the module will detect the structural analogues in MPDB.

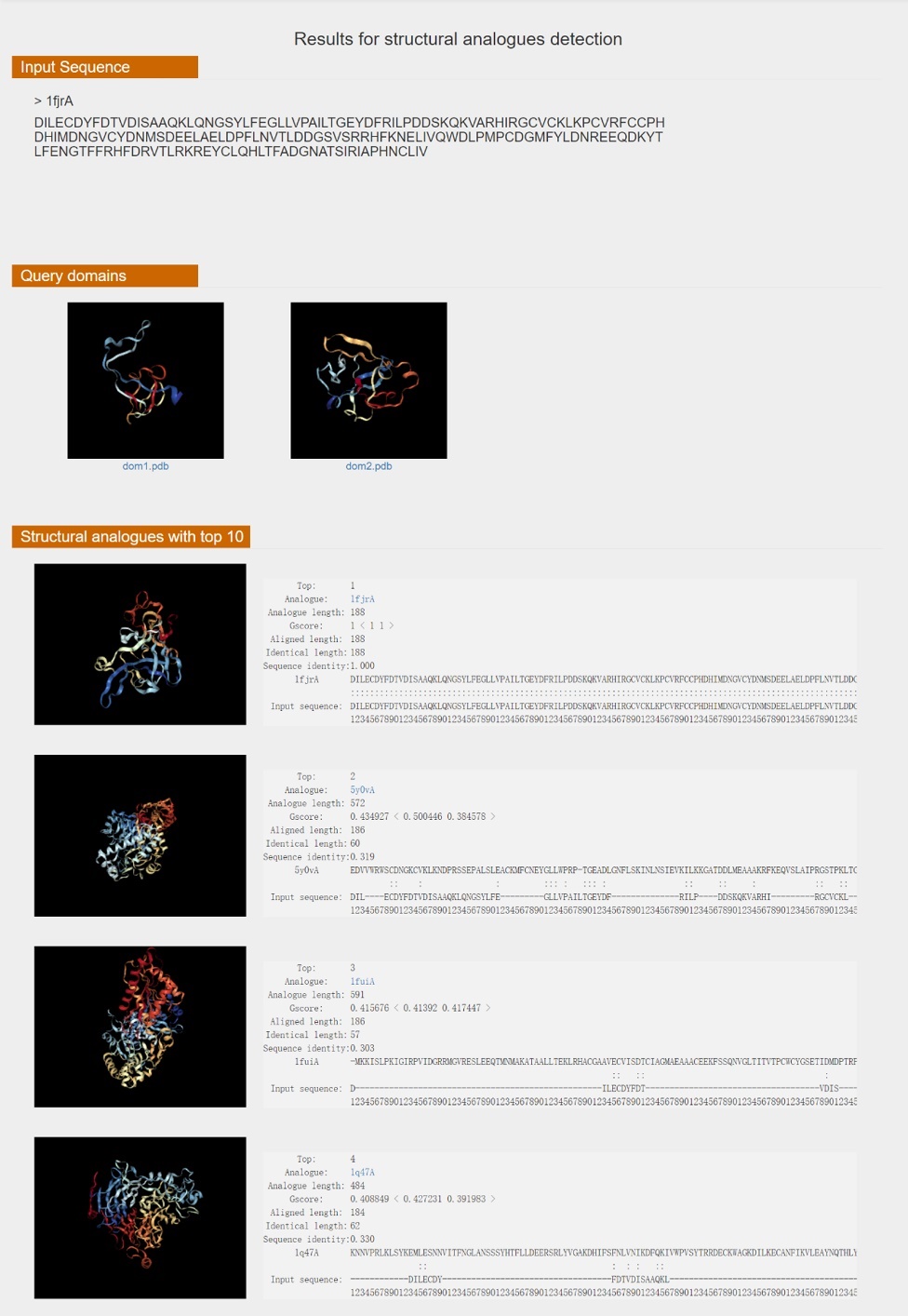

**Fig. S4.** The results page of structural analogue detection module in the MPDB web server. Upon job completion, the top 200 structural analogues with the highest *G*_score_ and the corresponding information are output. The results display page will display the input sequence, query domains and the top 10 structural analogues. The top 200 structural analogues are saved in a *.tar.gz file. The corresponding information is saved in a *.txt file, which includes the analogue name, length, *G*_score_, aligned score of each domain, aligned length between query sequence and sequence of analogue, identical length and sequence identity.

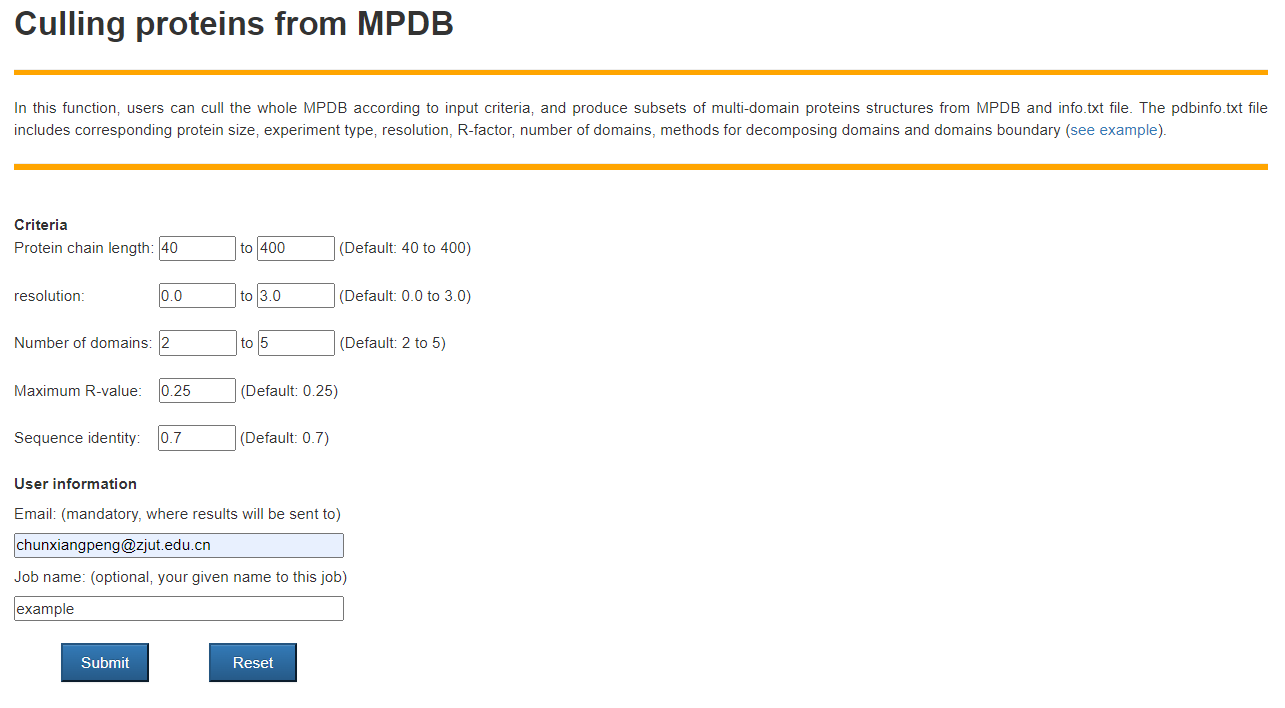

**Fig. S5.** The input page of culling module in the MPDB web server. According to the criteria input by the users, the culling module the module retrieves whole MPDB according to the criteria.

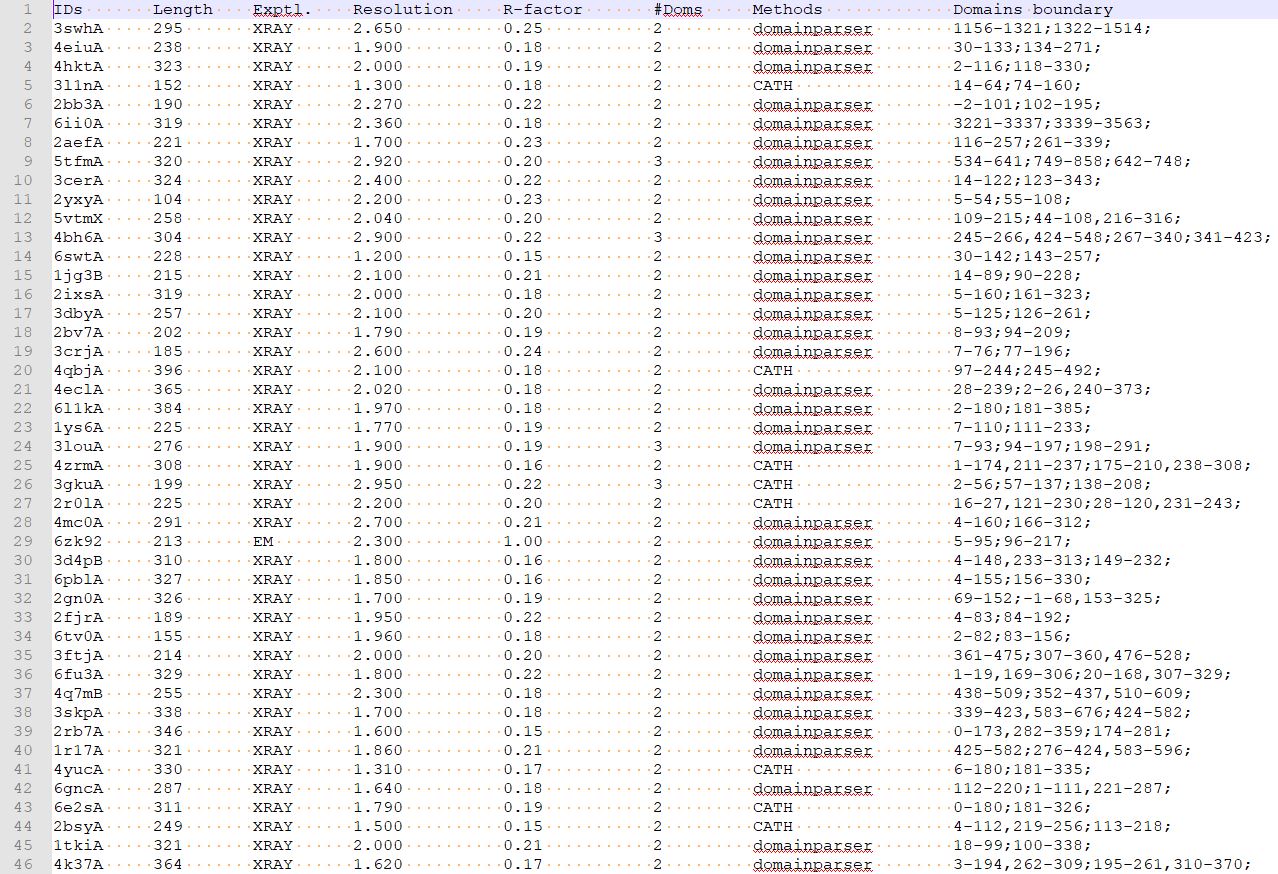

**Fig. S6.** The output results of culling module in the MPDB web server. After the retrieval is completed, the culling module will obtain the multi-domain protein structures that satisfy the criteria, and record the corresponding information, including the protein name, length, experimental measurement methods, resolution, R-factor, number of domains, and methods of defining domain boundary and domain boundary. The information is saved in a *.txt file. These multi-domain protein structures will be saved in a *.tar.gz file for users to download.

**Supplementary Tables**

**Table S1.** Protein length and number of domains on each test protein. #Domains represents the number of domains. 2dis represents the protein with 2 discontinuous domains.

| PDB ID | Length | #Domains | PDB ID | Length | #Domains | PDB ID | Length | #Domains |
| --- | --- | --- | --- | --- | --- | --- | --- | --- |
| 1cjyA  1efdN  1fjrA  1g87B  1hx6B  1iwaA  1m5qH  1mkfA  1mkmB  1nh2D  1pprM  1prrA  1q19A  1qwrA  1r71B  1rh1A  1rktA  1s6lA  1sp3A  1vz6A  1w3aA  1wv3A  1x7pA  1x9yA  1y11A  1yiqA  1zbuB  1ze1A  2ablA  2ahvA  2bkpA  2c1yA  2cxcA  2d1cA  2d7iA  2e9hA  2e9xB  2evrA  2ew9A  2fd5A  2gh8A  2gt1A  2gzaC  2hjqA  2hwjA  2ijd1  2iu7A  2iw2A  2jz4A  2kdyA  2kn4A  2mbgA  2nsfA  2nykA  2o6yA  2owbA  2qfiA  2qp2A  2qygA  2r5wB  2uu7A  2w4bA  2x7iA  2x8kC  2yilA  2yrqA  2zxcA  3a1iA  3a45A  3a56A  3ajvA  3aqkA  3arbA  3aujG  3b2zF  3b7wA  3bt3A  3c4tA  3craA  3d30A  3eo5A  3errA  3g79A  3h2tA  3hcsA  3hyiA  3i2dA  3iam2  3ifrA  3isqA  3j7aK  3k1rA  3k2iA  3kh5A  3kjpA  3kt1A  3ktmE  3kzwA  3l76A  3ld1A  3lsgA  3me4A  3ml4C  3mx2B  3mzfA  3njaB  3nqiA  3nt8A  3og5A  3oh0A  3pcsB  3po3S  3pxpA  3qavA  3qf4B  3qjjA  3qtdA  3r6bA  3rh7A | 630  262  188  613  396  441  123  371  247  102  312  173  500  321  116  490  207  181  436  365  312  181  265  346  356  684  289  308  163  512  202  231  138  495  536  157  175  240  149  180  544  323  336  99  184  644  159  479  299  261  158  265  240  235  514  262  286  513  429  345  370  455  296  243  131  173  643  508  288  291  174  390  274  139  293  543  129  241  239  212  169  527  475  326  157  286  279  179  483  384  129  193  411  279  293  557  171  494  585  359  101  169  203  517  362  107  240  401  155  593  355  164  288  219  588  243  435  115  289 | 2  2  2  2  2  2  2  2  2  2  2  2  2  2  2  2  2  2  2  2  2  2  2  2  2  2  2  2  2  2  2  2  2  2  2  2  2  2  2  2  2  2  2  2  2  2  2  2  2  2  2  2  2  2  2  2  2  2  2  2  2  2  2  2  2  2  2  2  2  2  2  2  2  2  2  2  2  2  2  2  2  2  2  2  2  2  2  2  2  2  2  2  2  2  2  2  2  2  2  2  2  2  2  2  2  2  2  2  2  2  2  2  2  2  2  2  2  2  2 | 3rwxA  3sb4A  3swjA  3t58B  3t7jA  3u07C  3u0oB  3u9gA  3ub1D  3uitD  3uo3A  3v7oB  3vr8B  3wkuA  3zvmA  4acoA  4ap5A  4axdA  4bfiB  4bt9B  4cczA  4d0nB  4d1iG  4dj3A  4dqaA  4eo3A  4eogA  4etxA  4fguA  4fkcA  4fxkC  4gbyA  4ggmX  4gslA  4gyjA  4h3tA  4hmoA  4ie6A  4l5gA  4lpqA  4m8rA  4n06B  4nj5A  4opaB  4qkuB  4up9A  4w7sA  1bf2A  1bhgA  1f7uA  1fx7A  1griA  1h88C  1m8pB  1ni5A  1q25A  1uzjA  1zpuA  1zy9A  2b5uA  2ewfA  2piaA  2r7dA  2uwnA  2v0nA  2vgmA  2wqrB  2y25B  2yk0A  2zzqA  3bt1U  3c1yA  3cw2C  3f83A  3fc3A  3gbgA  3h5cB  3ibjA  3ippB  3jymB  3kbgA  3mc8A  3npfA  3orjA  3plaA  3qe9Y  3qjoA  3qphA  3qyeA  3rimA  3rrpA  3soaA  3tixD  3tp9A  3ua3A  3uj0A  3vn4A  3vsmA  3w1bA  3zh9B  4alzA  4ax8A  4b3iA  4bd9B  4c0aB  4c0sA  4dimA  4indA  4jdzB  4kc3B  4kikB  4lmfA  4lziA  4m9pA  4pt5A  4uwhA  1c1zA  1d2pA  1k7tA | 239  323  246  509  237  382  335  217  251  257  184  203  249  405  385  452  373  437  202  228  611  371  544  298  358  328  460  300  407  370  290  475  270  598  618  518  310  404  162  206  398  340  482  197  427  514  444  750  611  606  230  211  152  571  428  428  162  504  522  465  574  321  452  187  459  354  322  314  698  469  273  349  251  508  190  260  263  661  420  307  187  257  306  415  375  345  491  335  317  697  455  436  424  467  648  502  375  638  589  339  212  456  736  165  355  449  382  383  447  278  648  276  278  292  319  553  325  373  170 | 2  2  2  2  2  2  2  2  2  2  2  2  2  2  2  2  2  2  2  2  2  2  2  2  2  2  2  2  2  2  2  2  2  2  2  2  2  2  2  2  2  2  2  2  2  2  2  3  3  3  3  3  3  3  3  3  3  3  3  3  3  3  3  3  3  3  3  3  3  3  3  3  3  3  3  3  3  3  3  3  3  3  3  3  3  3  3  3  3  3  3  3  3  3  3  3  3  3  3  3  3  3  3  3  3  3  3  3  3  3  3  3  3  3  3  3  5  4  4 | 1kfqA  1ldjA  1nyqB  1ug9A  1z1wA  2au3A  2ii2A  2olsA  2ra1A  2v5dA  2xt6A  2zpaB  3apoA  3b43A  3gf5B  3hjlA  3kq4B  3kwlA  3ob8A  3opfB  3p53A  3pvlA  3r05A  3ubhA  3w2wA  3zniA  4aimA  4ak1A  4aq1A  4fe9A  4h2aA  4i5sB  4iggB  4j9vA  4k3bA  4kwuA  4m00A  1bp1A  1cjsA  1ck1A  1ecrA  1f5qD  1fa9A  1gu7A  1itwA  1jkiA  1n80A  1nzjA  1qhdA  1qz9A  1sb7B  1vk1A  1vrmA  1xvuA  1yy3A  1z87A  2a1sC  2a3lA  2bt1A  2bydA  2c43A  2dfyC  2dlaA  2g3pA  2gg6A  2gsyE  2gzoA  2j2cA  2kfwA  2l9yA  2ntyB  2r3vA  2r58A  2w4mA  2x0cA  2y51A  2yb0E  2z86C  3afoA  3bu2A  3cvzA  3dupA  3eswA  3eukH  3fi7A  3fvvA  3gmsA  3hzzB  3m1uA  3mw8A  3mwcA  3nsjA  3ntkA  3oaaG  3ptyA  3rfyA  3seoB  3spgA  3u0kA  3vlaA  3vstA  4aqfB  4b21A  4dt4A  4dtfA  4ewtA  4f23A  4fzbC  4g1pA  4gfqA  4hvzA  4il6B  4jxkA  4m8mB  4mzyA  4onyA  4pyhA  4rg1A | 567  725  645  1019  780  403  304  725  413  722  1055  651  688  569  380  321  457  493  1024  493  494  598  1008  409  618  391  698  612  721  450  708  395  771  456  772  1030  495  456  212  239  305  248  833  364  740  525  328  273  397  404  340  223  304  402  315  263  393  616  254  283  280  158  222  201  445  433  187  470  196  167  337  390  212  250  308  529  264  603  360  189  248  279  332  459  177  223  331  447  410  233  388  530  169  284  284  356  226  332  396  428  638  474  207  160  359  389  509  207  479  186  214  505  327  574  495  583  293  285 | 4  7  4  4  4  4  4  4  5  4  4  4  7  6  7  4  4  4  5  4  4  5  7  4  4  4  4  6  6  4  4  4  6  4  6  6  4  2dis  2dis  2dis  2dis  2dis  2dis  2dis  2dis  2dis  2dis  2dis  2dis  2dis  2dis  2dis  2dis  2dis  2dis  2dis  2dis  2dis  2dis  2dis  2dis  2dis  2dis  2dis  2dis  2dis  2dis  2dis  2dis  2dis  2dis  2dis  2dis  2dis  2dis  2dis  2dis  2dis  2dis  2dis  2dis  2dis  2dis  2dis  2dis  2dis  2dis  2dis  2dis  2dis  2dis  2dis  2dis  2dis  2dis  2dis  2dis  2dis  2dis  2dis  2dis  2dis  2dis  2dis  2dis  2dis  2dis  2dis  2dis  2dis  2dis  2dis  2dis  2dis  2dis  2dis  2dis  2dis |

**Table S2.** Coverage scores of each test protein in MPDB and DEMO-lib under the sequence identity cutoff of 30%.

| PDB ID | Coverage score | | PDB ID | Coverage score | | PDB ID | Coverage score | |
| --- | --- | --- | --- | --- | --- | --- | --- | --- |
|  | MPDB | DEMO-lib |  | MPDB | DEMO-lib |  | MPDB | DEMO-lib |
| 1cjyA  1efdN  1fjrA  1g87B  1hx6B  1iwaA  1m5qH  1mkfA  1mkmB  1nh2D  1pprM  1prrA  1q19A  1qwrA  1r71B  1rh1A  1rktA  1s6lA  1sp3A  1vz6A  1w3aA  1wv3A  1x7pA  1x9yA  1y11A  1yiqA  1zbuB  1ze1A  2ablA  2ahvA  2bkpA  2c1yA  2cxcA  2d1cA  2d7iA  2e9hA  2e9xB  2evrA  2ew9A  2fd5A  2gh8A  2gt1A  2gzaC  2hjqA  2hwjA  2ijd1  2iu7A  2iw2A  2jz4A  2kdyA  2kn4A  2mbgA  2nsfA  2nykA  2o6yA  2owbA  2qfiA  2qp2A  2qygA  2r5wB  2uu7A  2w4bA  2x7iA  2x8kC  2yilA  2yrqA  2zxcA  3a1iA  3a45A  3a56A  3ajvA  3aqkA  3arbA  3aujG  3b2zF  3b7wA  3bt3A  3c4tA  3craA  3d30A  3eo5A  3errA  3g79A  3h2tA  3hcsA  3hyiA  3i2dA  3iam2  3ifrA  3isqA  3j7aK  3k1rA  3k2iA  3kh5A  3kjpA  3kt1A  3ktmE  3kzwA  3l76A  3ld1A  3lsgA  3me4A  3ml4C  3mx2B  3mzfA  3njaB  3nqiA  3nt8A  3og5A  3oh0A  3pcsB  3po3S  3pxpA  3qavA  3qf4B  3qjjA  3qtdA  3r6bA  3rh7A | 0.38  0.83  0.45  0.61  0.67  0.84  0.48  0.43  0.65  0.49  0.45  0.47  0.56  0.83  0.62  0.40  0.71  0.44  0.39  0.43  0.48  0.49  0.64  0.47  0.49  0.49  0.55  0.62  0.55  0.74  0.50  0.56  0.63  0.65  0.45  0.47  0.63  0.58  0.55  0.78  0.52  0.68  0.48  0.50  0.47  0.67  0.50  0.80  0.48  0.52  0.48  0.63  0.47  0.67  0.72  0.93  0.44  0.50  0.89  0.46  0.77  0.36  0.77  0.51  0.54  0.44  0.33  0.55  0.80  0.43  0.68  0.74  0.84  0.49  0.63  0.93  0.74  0.58  0.45  0.51  0.43  0.69  0.79  0.39  0.47  0.50  0.50  0.55  0.91  0.71  0.84  0.46  0.59  0.83  0.52  0.43  0.45  0.86  0.52  0.42  0.77  0.56  0.49  0.44  0.76  0.48  0.56  0.40  0.59  0.65  0.47  0.45  0.42  0.81  0.80  0.70  0.42  0.48  0.45 | 0.36  0.80  0.43  0.55  0.60  0.79  0.47  0.42  0.57  0.49  0.43  0.45  0.46  0.67  0.52  0.37  0.70  0.43  0.38  0.42  0.47  0.48  0.54  0.46  0.45  0.47  0.55  0.55  0.52  0.67  0.49  0.50  0.56  0.65  0.42  0.47  0.55  0.54  0.54  0.78  0.46  0.66  0.47  0.49  0.46  0.62  0.48  0.75  0.46  0.46  0.48  0.62  0.45  0.66  0.52  0.90  0.42  0.46  0.87  0.43  0.70  0.36  0.75  0.49  0.48  0.43  0.32  0.48  0.77  0.41  0.57  0.66  0.82  0.47  0.55  0.85  0.72  0.50  0.43  0.50  0.41  0.61  0.71  0.38  0.47  0.49  0.46  0.46  0.90  0.71  0.63  0.45  0.58  0.57  0.43  0.42  0.44  0.76  0.48  0.41  0.74  0.51  0.48  0.41  0.65  0.46  0.55  0.39  0.53  0.61  0.41  0.44  0.41  0.81  0.71  0.64  0.42  0.47  0.44 | 3rwxA  3sb4A  3swjA  3t58B  3t7jA  3u07C  3u0oB  3u9gA  3ub1D  3uitD  3uo3A  3v7oB  3vr8B  3wkuA  3zvmA  4acoA  4ap5A  4axdA  4bfiB  4bt9B  4cczA  4d0nB  4d1iG  4dj3A  4dqaA  4eo3A  4eogA  4etxA  4fguA  4fkcA  4fxkC  4gbyA  4ggmX  4gslA  4gyjA  4h3tA  4hmoA  4ie6A  4l5gA  4lpqA  4m8rA  4n06B  4nj5A  4opaB  4qkuB  4up9A  4w7sA  1bf2A  1bhgA  1f7uA  1fx7A  1griA  1h88C  1m8pB  1ni5A  1q25A  1uzjA  1zpuA  1zy9A  2b5uA  2ewfA  2piaA  2r7dA  2uwnA  2v0nA  2vgmA  2wqrB  2y25B  2yk0A  2zzqA  3bt1U  3c1yA  3cw2C  3f83A  3fc3A  3gbgA  3h5cB  3ibjA  3ippB  3jymB  3kbgA  3mc8A  3npfA  3orjA  3plaA  3qe9Y  3qjoA  3qphA  3qyeA  3rimA  3rrpA  3soaA  3tixD  3tp9A  3ua3A  3uj0A  3vn4A  3vsmA  3w1bA  3zh9B  4alzA  4ax8A  4b3iA  4bd9B  4c0aB  4c0sA  4dimA  4indA  4jdzB  4kc3B  4kikB  4lmfA  4lziA  4m9pA  4pt5A  4uwhA  1c1zA  1d2pA  1k7tA | 0.45  0.53  0.51  0.42  0.65  0.45  0.73  0.44  0.41  0.48  0.61  0.46  0.52  0.42  0.42  0.41  0.61  0.40  0.79  0.52  0.50  0.76  0.65  0.56  0.45  0.48  0.60  0.43  0.43  0.88  0.55  0.78  0.44  0.40  0.80  0.54  0.74  0.42  0.50  0.52  0.54  0.75  0.37  0.45  0.57  0.73  0.53  0.75  0.83  0.70  0.65  0.46  0.49  0.59  0.46  0.40  0.42  0.87  0.78  0.37  0.39  0.63  0.70  0.48  0.41  0.56  0.58  0.44  0.42  0.44  0.42  0.43  0.55  0.42  0.44  0.43  0.80  0.44  0.60  0.43  0.42  0.52  0.56  0.47  0.45  0.72  0.55  0.42  0.75  0.82  0.80  0.63  0.38  0.46  0.42  0.70  0.39  0.76  0.64  0.52  0.46  0.45  0.60  0.43  0.50  0.72  0.80  0.41  0.60  0.72  0.63  0.45  0.41  0.44  0.65  0.85  0.44  0.39  0.46 | 0.44  0.51  0.48  0.38  0.63  0.40  0.71  0.43  0.40  0.43  0.55  0.46  0.43  0.40  0.41  0.37  0.54  0.36  0.77  0.50  0.47  0.70  0.64  0.46  0.43  0.46  0.42  0.42  0.42  0.87  0.47  0.73  0.44  0.38  0.78  0.45  0.74  0.40  0.48  0.51  0.53  0.68  0.36  0.44  0.45  0.72  0.50  0.73  0.80  0.64  0.50  0.43  0.48  0.47  0.43  0.38  0.42  0.74  0.74  0.36  0.37  0.63  0.46  0.47  0.39  0.51  0.52  0.42  0.39  0.42  0.42  0.42  0.51  0.41  0.43  0.42  0.78  0.42  0.56  0.42  0.42  0.50  0.49  0.41  0.40  0.65  0.42  0.42  0.70  0.60  0.77  0.59  0.37  0.43  0.42  0.48  0.38  0.71  0.52  0.49  0.46  0.43  0.53  0.43  0.45  0.51  0.78  0.39  0.58  0.71  0.42  0.44  0.41  0.44  0.57  0.60  0.41  0.38  0.46 | 1kfqA  1ldjA  1nyqB  1ug9A  1z1wA  2au3A  2ii2A  2olsA  2ra1A  2v5dA  2xt6A  2zpaB  3apoA  3b43A  3gf5B  3hjlA  3kq4B  3kwlA  3ob8A  3opfB  3p53A  3pvlA  3r05A  3ubhA  3w2wA  3zniA  4aimA  4ak1A  4aq1A  4fe9A  4h2aA  4i5sB  4iggB  4j9vA  4k3bA  4kwuA  4m00A  1bp1A  1cjsA  1ck1A  1ecrA  1f5qD  1fa9A  1gu7A  1itwA  1jkiA  1n80A  1nzjA  1qhdA  1qz9A  1sb7B  1vk1A  1vrmA  1xvuA  1yy3A  1z87A  2a1sC  2a3lA  2bt1A  2bydA  2c43A  2dfyC  2dlaA  2g3pA  2gg6A  2gsyE  2gzoA  2j2cA  2kfwA  2l9yA  2ntyB  2r3vA  2r58A  2w4mA  2x0cA  2y51A  2yb0E  2z86C  3afoA  3bu2A  3cvzA  3dupA  3eswA  3eukH  3fi7A  3fvvA  3gmsA  3hzzB  3m1uA  3mw8A  3mwcA  3nsjA  3ntkA  3oaaG  3ptyA  3rfyA  3seoB  3spgA  3u0kA  3vlaA  3vstA  4aqfB  4b21A  4dt4A  4dtfA  4ewtA  4f23A  4fzbC  4g1pA  4gfqA  4hvzA  4il6B  4jxkA  4m8mB  4mzyA  4onyA  4pyhA  4rg1A | 0.73  0.44  0.59  0.53  0.89  0.65  0.84  0.41  0.41  0.62  0.50  0.37  0.37  0.46  0.39  0.43  0.41  0.41  0.52  0.37  0.40  0.60  0.33  0.48  0.57  0.40  0.36  0.38  0.34  0.39  0.40  0.46  0.55  0.58  0.38  0.67  0.38  0.44  0.65  0.77  0.38  0.80  0.50  0.77  0.48  0.58  0.37  0.74  0.43  0.85  0.45  0.48  0.46  0.86  0.41  0.38  0.41  0.44  0.50  0.51  0.51  0.43  0.51  0.40  0.87  0.45  0.42  0.43  0.43  0.41  0.41  0.70  0.42  0.74  0.45  0.85  0.46  0.37  0.69  0.43  0.40  0.49  0.50  0.59  0.61  0.76  0.88  0.72  0.48  0.62  0.86  0.50  0.82  0.64  0.45  0.38  0.45  0.75  0.55  0.72  0.49  0.39  0.84  0.58  0.40  0.79  0.60  0.61  0.74  0.51  0.46  0.54  0.84  0.36  0.56  0.73  0.58  0.54 | 0.73  0.43  0.56  0.51  0.85  0.54  0.82  0.38  0.39  0.54  0.48  0.36  0.34  0.44  0.38  0.41  0.40  0.39  0.51  0.39  0.39  0.52  0.31  0.48  0.39  0.40  0.34  0.37  0.33  0.38  0.36  0.43  0.36  0.54  0.33  0.66  0.37  0.41  0.63  0.71  0.38  0.77  0.46  0.76  0.46  0.49  0.36  0.74  0.39  0.82  0.44  0.41  0.41  0.70  0.40  0.37  0.40  0.44  0.43  0.45  0.45  0.42  0.49  0.40  0.82  0.42  0.40  0.40  0.43  0.41  0.41  0.68  0.41  0.72  0.43  0.84  0.45  0.36  0.61  0.42  0.40  0.47  0.42  0.57  0.55  0.70  0.88  0.71  0.40  0.59  0.86  0.47  0.63  0.53  0.44  0.38  0.44  0.46  0.41  0.68  0.47  0.38  0.79  0.55  0.39  0.78  0.50  0.45  0.72  0.49  0.43  0.45  0.84  0.34  0.58  0.71  0.54  0.50 |

**Table S3.** Under the sequence identity cutoff of 30%, average coverage scores of 2dom proteins, 3dom proteins, m4dom proteins, 2dis proteins and all test protein in MPDB and DEMO-lib. *P*-value is determined by Student’s t-tests.

| Method | 2dom (166) | 3dom (69) | m4dom (40) | 2dis (81) | Total (356) | *P*-value |
| --- | --- | --- | --- | --- | --- | --- |
| MPDB | 0.57 | 0.56 | 0.48 | 0.56 | 0.56 | NA |
| DEMO-lib | 0.53 | 0.50 | 0.45 | 0.53 | 0.52 | 2.04E-04 |

**Table S4.** Coverage scores of each test protein in MPDB and DEMO-lib under the sequence identity cutoff of 50%.

| PDB ID | Coverage score | | PDB ID | Coverage score | | PDB ID | Coverage score | |
| --- | --- | --- | --- | --- | --- | --- | --- | --- |
|  | MPDB | DEMO-lib |  | MPDB | DEMO-lib |  | MPDB | DEMO-lib |
| 1cjyA  1efdN  1fjrA  1g87B  1hx6B  1iwaA  1m5qH  1mkfA  1mkmB  1nh2D  1pprM  1prrA  1q19A  1qwrA  1r71B  1rh1A  1rktA  1s6lA  1sp3A  1vz6A  1w3aA  1wv3A  1x7pA  1x9yA  1y11A  1yiqA  1zbuB  1ze1A  2ablA  2ahvA  2bkpA  2c1yA  2cxcA  2d1cA  2d7iA  2e9hA  2e9xB  2evrA  2ew9A  2fd5A  2gh8A  2gt1A  2gzaC  2hjqA  2hwjA  2ijd1  2iu7A  2iw2A  2jz4A  2kdyA  2kn4A  2mbgA  2nsfA  2nykA  2o6yA  2owbA  2qfiA  2qp2A  2qygA  2r5wB  2uu7A  2w4bA  2x7iA  2x8kC  2yilA  2yrqA  2zxcA  3a1iA  3a45A  3a56A  3ajvA  3aqkA  3arbA  3aujG  3b2zF  3b7wA  3bt3A  3c4tA  3craA  3d30A  3eo5A  3errA  3g79A  3h2tA  3hcsA  3hyiA  3i2dA  3iam2  3ifrA  3isqA  3j7aK  3k1rA  3k2iA  3kh5A  3kjpA  3kt1A  3ktmE  3kzwA  3l76A  3ld1A  3lsgA  3me4A  3ml4C  3mx2B  3mzfA  3njaB  3nqiA  3nt8A  3og5A  3oh0A  3pcsB  3po3S  3pxpA  3qavA  3qf4B  3qjjA  3qtdA  3r6bA  3rh7A | 0.47  0.85  0.45  0.71  0.67  0.92  0.51  0.43  0.65  0.64  0.50  0.49  0.59  0.83  0.63  0.41  0.71  0.45  0.39  0.44  0.49  0.50  0.70  0.47  0.49  0.62  0.56  0.78  0.59  0.74  0.50  0.57  0.70  0.65  0.69  0.49  0.63  0.66  0.55  0.78  0.56  0.68  0.48  0.52  0.47  0.70  0.57  0.82  0.49  0.52  0.49  0.63  0.49  0.67  0.93  0.94  0.49  0.50  0.89  0.55  0.79  0.51  0.79  0.53  0.56  0.48  0.45  0.63  0.80  0.43  0.73  0.74  0.86  0.51  0.69  0.93  0.74  0.59  0.47  0.61  0.44  0.70  0.79  0.41  0.48  0.50  0.55  0.78  0.92  0.85  0.87  0.46  0.59  0.83  0.52  0.43  0.45  0.89  0.57  0.43  0.85  0.63  0.50  0.45  0.78  0.50  0.56  0.41  0.73  0.65  0.47  0.50  0.42  0.81  0.88  0.73  0.44  0.60  0.47 | 0.41  0.83  0.43  0.57  0.60  0.89  0.49  0.42  0.58  0.59  0.48  0.48  0.48  0.67  0.55  0.39  0.70  0.45  0.38  0.43  0.47  0.49  0.61  0.46  0.46  0.58  0.56  0.62  0.56  0.67  0.50  0.52  0.61  0.65  0.50  0.48  0.56  0.56  0.54  0.78  0.46  0.66  0.48  0.50  0.46  0.66  0.49  0.76  0.47  0.48  0.48  0.62  0.47  0.66  0.81  0.93  0.46  0.46  0.88  0.48  0.77  0.41  0.76  0.50  0.51  0.46  0.45  0.56  0.77  0.42  0.65  0.67  0.83  0.50  0.62  0.86  0.72  0.53  0.46  0.57  0.42  0.62  0.75  0.39  0.48  0.49  0.47  0.47  0.91  0.79  0.72  0.46  0.58  0.57  0.43  0.42  0.44  0.88  0.50  0.42  0.79  0.57  0.50  0.41  0.69  0.49  0.56  0.39  0.63  0.61  0.41  0.48  0.42  0.81  0.85  0.64  0.43  0.52  0.44 | 3rwxA  3sb4A  3swjA  3t58B  3t7jA  3u07C  3u0oB  3u9gA  3ub1D  3uitD  3uo3A  3v7oB  3vr8B  3wkuA  3zvmA  4acoA  4ap5A  4axdA  4bfiB  4bt9B  4cczA  4d0nB  4d1iG  4dj3A  4dqaA  4eo3A  4eogA  4etxA  4fguA  4fkcA  4fxkC  4gbyA  4ggmX  4gslA  4gyjA  4h3tA  4hmoA  4ie6A  4l5gA  4lpqA  4m8rA  4n06B  4nj5A  4opaB  4qkuB  4up9A  4w7sA  1bf2A  1bhgA  1f7uA  1fx7A  1griA  1h88C  1m8pB  1ni5A  1q25A  1uzjA  1zpuA  1zy9A  2b5uA  2ewfA  2piaA  2r7dA  2uwnA  2v0nA  2vgmA  2wqrB  2y25B  2yk0A  2zzqA  3bt1U  3c1yA  3cw2C  3f83A  3fc3A  3gbgA  3h5cB  3ibjA  3ippB  3jymB  3kbgA  3mc8A  3npfA  3orjA  3plaA  3qe9Y  3qjoA  3qphA  3qyeA  3rimA  3rrpA  3soaA  3tixD  3tp9A  3ua3A  3uj0A  3vn4A  3vsmA  3w1bA  3zh9B  4alzA  4ax8A  4b3iA  4bd9B  4c0aB  4c0sA  4dimA  4indA  4jdzB  4kc3B  4kikB  4lmfA  4lziA  4m9pA  4pt5A  4uwhA  1c1zA  1d2pA  1k7tA | 0.45  0.54  0.51  0.47  0.65  0.45  0.79  0.44  0.41  0.48  0.61  0.46  0.71  0.43  0.42  0.42  0.69  0.44  0.79  0.54  0.50  0.77  0.65  0.60  0.47  0.48  0.60  0.44  0.90  0.89  0.72  0.81  0.46  0.41  0.80  0.71  0.74  0.42  0.50  0.52  0.54  0.76  0.51  0.46  0.57  0.88  0.55  0.79  0.92  0.83  0.66  0.52  0.57  0.69  0.47  0.40  0.48  0.90  0.78  0.37  0.39  0.64  0.84  0.51  0.42  0.57  0.58  0.45  0.42  0.44  0.42  0.43  0.57  0.45  0.45  0.44  0.80  0.45  0.86  0.44  0.86  0.52  0.56  0.47  0.54  0.72  0.71  0.42  0.78  0.93  0.93  0.65  0.41  0.48  0.44  0.70  0.41  0.76  0.64  0.53  0.47  0.45  0.81  0.44  0.52  0.74  0.80  0.43  0.60  0.72  0.63  0.63  0.42  0.48  0.86  0.85  0.45  0.39  0.50 | 0.44  0.52  0.48  0.38  0.63  0.41  0.74  0.44  0.40  0.44  0.56  0.46  0.53  0.41  0.41  0.37  0.63  0.40  0.77  0.52  0.47  0.72  0.64  0.46  0.46  0.46  0.42  0.42  0.47  0.89  0.58  0.75  0.44  0.38  0.79  0.59  0.74  0.40  0.50  0.51  0.53  0.68  0.41  0.45  0.45  0.83  0.53  0.77  0.86  0.70  0.51  0.48  0.54  0.56  0.44  0.38  0.46  0.85  0.74  0.36  0.37  0.63  0.60  0.49  0.40  0.52  0.56  0.44  0.39  0.42  0.42  0.42  0.55  0.44  0.44  0.42  0.78  0.42  0.56  0.42  0.73  0.50  0.49  0.41  0.43  0.66  0.55  0.42  0.76  0.92  0.85  0.62  0.40  0.46  0.43  0.49  0.39  0.71  0.52  0.50  0.47  0.43  0.58  0.43  0.46  0.56  0.79  0.41  0.58  0.71  0.42  0.55  0.42  0.46  0.63  0.77  0.42  0.39  0.48 | 1kfqA  1ldjA  1nyqB  1ug9A  1z1wA  2au3A  2ii2A  2olsA  2ra1A  2v5dA  2xt6A  2zpaB  3apoA  3b43A  3gf5B  3hjlA  3kq4B  3kwlA  3ob8A  3opfB  3p53A  3pvlA  3r05A  3ubhA  3w2wA  3zniA  4aimA  4ak1A  4aq1A  4fe9A  4h2aA  4i5sB  4iggB  4j9vA  4k3bA  4kwuA  4m00A  1bp1A  1cjsA  1ck1A  1ecrA  1f5qD  1fa9A  1gu7A  1itwA  1jkiA  1n80A  1nzjA  1qhdA  1qz9A  1sb7B  1vk1A  1vrmA  1xvuA  1yy3A  1z87A  2a1sC  2a3lA  2bt1A  2bydA  2c43A  2dfyC  2dlaA  2g3pA  2gg6A  2gsyE  2gzoA  2j2cA  2kfwA  2l9yA  2ntyB  2r3vA  2r58A  2w4mA  2x0cA  2y51A  2yb0E  2z86C  3afoA  3bu2A  3cvzA  3dupA  3eswA  3eukH  3fi7A  3fvvA  3gmsA  3hzzB  3m1uA  3mw8A  3mwcA  3nsjA  3ntkA  3oaaG  3ptyA  3rfyA  3seoB  3spgA  3u0kA  3vlaA  3vstA  4aqfB  4b21A  4dt4A  4dtfA  4ewtA  4f23A  4fzbC  4g1pA  4gfqA  4hvzA  4il6B  4jxkA  4m8mB  4mzyA  4onyA  4pyhA  4rg1A | 0.80  0.60  0.62  0.53  0.90  0.65  0.84  0.42  0.42  0.73  0.62  0.42  0.37  0.46  0.39  0.44  0.41  0.41  0.89  0.42  0.40  0.63  0.33  0.48  0.57  0.53  0.73  0.38  0.34  0.39  0.48  0.49  0.56  0.58  0.49  0.67  0.39  0.48  0.82  0.88  0.40  0.80  0.88  0.79  0.48  0.63  0.37  0.88  0.43  0.85  0.50  0.48  0.65  0.86  0.52  0.40  0.41  0.44  0.50  0.51  0.51  0.50  0.51  0.41  0.90  0.56  0.49  0.47  0.44  0.43  0.42  0.70  0.87  0.74  0.47  0.86  0.54  0.37  0.70  0.50  0.43  0.49  0.53  0.59  0.63  0.77  0.88  0.76  0.48  0.64  0.89  0.50  0.82  0.76  0.68  0.39  0.46  0.76  0.56  0.78  0.49  0.43  0.85  0.60  0.40  0.81  0.89  0.69  0.78  0.67  0.46  0.54  0.90  0.73  0.63  0.75  0.58  0.54 | 0.75  0.47  0.60  0.51  0.88  0.54  0.82  0.38  0.41  0.58  0.50  0.36  0.35  0.44  0.39  0.43  0.40  0.40  0.66  0.43  0.39  0.56  0.31  0.48  0.39  0.43  0.47  0.38  0.33  0.38  0.37  0.47  0.36  0.54  0.36  0.66  0.37  0.45  0.78  0.77  0.38  0.77  0.70  0.77  0.46  0.54  0.36  0.83  0.40  0.82  0.49  0.42  0.51  0.70  0.44  0.40  0.40  0.44  0.44  0.46  0.45  0.46  0.50  0.40  0.86  0.48  0.42  0.49  0.43  0.41  0.41  0.68  0.59  0.72  0.44  0.84  0.47  0.37  0.63  0.47  0.41  0.47  0.46  0.57  0.58  0.72  0.88  0.74  0.41  0.63  0.88  0.47  0.71  0.57  0.45  0.38  0.45  0.50  0.42  0.75  0.47  0.42  0.81  0.57  0.39  0.80  0.80  0.50  0.75  0.59  0.44  0.45  0.86  0.58  0.65  0.74  0.54  0.50 |

**Table S5.** Under the sequence identity cutoff of 50%, average coverage scores of 2dom proteins, 3dom proteins, m4dom proteins, 2dis proteins and all test protein in MPDB and DEMO-lib. *P*-value is determined by Student’s t-tests.

| Method | 2dom (166) | 3dom (69) | m4dom (40) | 2dis (81) | Total (356) | *P*-value |
| --- | --- | --- | --- | --- | --- | --- |
| MPDB | 0.60 | 0.60 | 0.53 | 0.62 | 0.60 | NA |
| DEMO-lib | 0.56 | 0.54 | 0.47 | 0.56 | 0.55 | 5.99E-06 |

**Table S6.** Coverage scores of each test protein in MPDB and DEMO-lib under the sequence identity cutoff of 70%.

| PDB ID | Coverage score | | PDB ID | Coverage score | | PDB ID | Coverage score | |
| --- | --- | --- | --- | --- | --- | --- | --- | --- |
|  | MPDB | DEMO-lib |  | MPDB | DEMO-lib |  | MPDB | DEMO-lib |
| 1cjyA  1efdN  1fjrA  1g87B  1hx6B  1iwaA  1m5qH  1mkfA  1mkmB  1nh2D  1pprM  1prrA  1q19A  1qwrA  1r71B  1rh1A  1rktA  1s6lA  1sp3A  1vz6A  1w3aA  1wv3A  1x7pA  1x9yA  1y11A  1yiqA  1zbuB  1ze1A  2ablA  2ahvA  2bkpA  2c1yA  2cxcA  2d1cA  2d7iA  2e9hA  2e9xB  2evrA  2ew9A  2fd5A  2gh8A  2gt1A  2gzaC  2hjqA  2hwjA  2ijd1  2iu7A  2iw2A  2jz4A  2kdyA  2kn4A  2mbgA  2nsfA  2nykA  2o6yA  2owbA  2qfiA  2qp2A  2qygA  2r5wB  2uu7A  2w4bA  2x7iA  2x8kC  2yilA  2yrqA  2zxcA  3a1iA  3a45A  3a56A  3ajvA  3aqkA  3arbA  3aujG  3b2zF  3b7wA  3bt3A  3c4tA  3craA  3d30A  3eo5A  3errA  3g79A  3h2tA  3hcsA  3hyiA  3i2dA  3iam2  3ifrA  3isqA  3j7aK  3k1rA  3k2iA  3kh5A  3kjpA  3kt1A  3ktmE  3kzwA  3l76A  3ld1A  3lsgA  3me4A  3ml4C  3mx2B  3mzfA  3njaB  3nqiA  3nt8A  3og5A  3oh0A  3pcsB  3po3S  3pxpA  3qavA  3qf4B  3qjjA  3qtdA  3r6bA  3rh7A | 0.47  0.85  0.45  0.78  0.67  0.99  0.51  0.43  0.65  0.64  0.50  0.49  0.59  0.83  0.63  0.41  0.71  0.45  0.39  0.46  0.49  0.50  0.70  0.47  0.49  0.67  0.56  0.78  0.59  0.74  0.50  0.57  0.70  0.65  0.69  0.49  0.63  0.66  0.55  0.78  0.69  0.68  0.48  0.52  0.47  0.71  0.72  0.82  0.49  0.56  0.49  0.63  0.49  0.67  0.93  0.94  0.49  0.50  0.92  0.55  0.84  0.51  0.79  0.53  0.56  0.49  0.45  0.63  0.80  0.43  0.73  0.74  0.96  0.56  0.78  0.93  0.74  0.59  0.47  0.61  0.44  0.70  0.79  0.41  0.52  0.50  0.55  0.78  0.92  0.85  0.92  0.46  0.66  0.83  0.52  0.43  0.45  0.89  0.57  0.43  0.85  0.63  0.50  0.45  0.83  0.50  0.56  0.41  0.76  0.65  0.47  0.50  0.42  0.81  0.88  0.73  0.44  0.63  0.47 | 0.41  0.83  0.43  0.62  0.60  0.90  0.49  0.42  0.58  0.59  0.48  0.48  0.48  0.67  0.55  0.39  0.70  0.45  0.38  0.43  0.47  0.49  0.61  0.46  0.46  0.62  0.56  0.62  0.56  0.67  0.50  0.52  0.61  0.65  0.50  0.48  0.56  0.56  0.54  0.78  0.51  0.66  0.48  0.50  0.46  0.66  0.54  0.76  0.47  0.52  0.48  0.62  0.47  0.66  0.81  0.94  0.46  0.46  0.89  0.48  0.81  0.41  0.76  0.50  0.51  0.47  0.45  0.56  0.77  0.42  0.65  0.67  0.92  0.55  0.66  0.86  0.72  0.53  0.46  0.57  0.42  0.65  0.75  0.39  0.48  0.49  0.47  0.47  0.91  0.79  0.76  0.46  0.62  0.57  0.43  0.42  0.44  0.88  0.50  0.42  0.79  0.57  0.50  0.41  0.76  0.49  0.56  0.39  0.68  0.61  0.41  0.48  0.42  0.81  0.85  0.64  0.43  0.54  0.44 | 3rwxA  3sb4A  3swjA  3t58B  3t7jA  3u07C  3u0oB  3u9gA  3ub1D  3uitD  3uo3A  3v7oB  3vr8B  3wkuA  3zvmA  4acoA  4ap5A  4axdA  4bfiB  4bt9B  4cczA  4d0nB  4d1iG  4dj3A  4dqaA  4eo3A  4eogA  4etxA  4fguA  4fkcA  4fxkC  4gbyA  4ggmX  4gslA  4gyjA  4h3tA  4hmoA  4ie6A  4l5gA  4lpqA  4m8rA  4n06B  4nj5A  4opaB  4qkuB  4up9A  4w7sA  1bf2A  1bhgA  1f7uA  1fx7A  1griA  1h88C  1m8pB  1ni5A  1q25A  1uzjA  1zpuA  1zy9A  2b5uA  2ewfA  2piaA  2r7dA  2uwnA  2v0nA  2vgmA  2wqrB  2y25B  2yk0A  2zzqA  3bt1U  3c1yA  3cw2C  3f83A  3fc3A  3gbgA  3h5cB  3ibjA  3ippB  3jymB  3kbgA  3mc8A  3npfA  3orjA  3plaA  3qe9Y  3qjoA  3qphA  3qyeA  3rimA  3rrpA  3soaA  3tixD  3tp9A  3ua3A  3uj0A  3vn4A  3vsmA  3w1bA  3zh9B  4alzA  4ax8A  4b3iA  4bd9B  4c0aB  4c0sA  4dimA  4indA  4jdzB  4kc3B  4kikB  4lmfA  4lziA  4m9pA  4pt5A  4uwhA  1c1zA  1d2pA  1k7tA | 0.45  0.54  0.55  0.47  0.65  0.45  0.79  0.44  0.41  0.48  0.61  0.46  0.86  0.43  0.42  0.42  0.69  0.44  0.79  0.54  0.50  0.77  0.67  0.69  0.47  0.48  0.60  0.44  0.90  0.90  0.72  0.81  0.46  0.41  0.80  0.71  0.74  0.42  0.50  0.52  0.54  0.76  0.51  0.46  0.64  0.88  0.55  0.79  0.92  0.83  0.80  0.52  0.60  0.73  0.47  0.40  0.48  0.90  0.78  0.37  0.39  0.64  0.84  0.53  0.42  0.57  0.62  0.47  0.44  0.44  0.51  0.43  0.57  0.48  0.45  0.44  0.80  0.45  0.86  0.44  0.86  0.56  0.56  0.47  0.56  0.72  0.71  0.45  0.79  0.93  0.93  0.66  0.41  0.48  0.44  0.70  0.41  0.76  0.64  0.53  0.47  0.45  0.81  0.44  0.52  0.76  0.80  0.43  0.61  0.75  0.67  0.64  0.42  0.48  0.86  0.86  0.45  0.41  0.50 | 0.44  0.52  0.53  0.38  0.63  0.41  0.74  0.44  0.40  0.44  0.56  0.46  0.63  0.41  0.41  0.37  0.63  0.40  0.77  0.52  0.47  0.72  0.67  0.55  0.46  0.46  0.42  0.42  0.47  0.89  0.58  0.75  0.44  0.38  0.79  0.59  0.74  0.40  0.50  0.51  0.53  0.68  0.41  0.45  0.51  0.87  0.53  0.77  0.86  0.70  0.59  0.48  0.54  0.59  0.44  0.38  0.46  0.85  0.74  0.36  0.37  0.63  0.60  0.49  0.40  0.52  0.57  0.45  0.42  0.42  0.51  0.42  0.55  0.46  0.44  0.42  0.78  0.42  0.56  0.42  0.73  0.55  0.49  0.41  0.45  0.66  0.55  0.44  0.78  0.92  0.89  0.63  0.40  0.46  0.43  0.50  0.39  0.71  0.52  0.50  0.47  0.43  0.58  0.43  0.46  0.64  0.79  0.41  0.59  0.71  0.51  0.56  0.42  0.46  0.63  0.81  0.42  0.39  0.48 | 1kfqA  1ldjA  1nyqB  1ug9A  1z1wA  2au3A  2ii2A  2olsA  2ra1A  2v5dA  2xt6A  2zpaB  3apoA  3b43A  3gf5B  3hjlA  3kq4B  3kwlA  3ob8A  3opfB  3p53A  3pvlA  3r05A  3ubhA  3w2wA  3zniA  4aimA  4ak1A  4aq1A  4fe9A  4h2aA  4i5sB  4iggB  4j9vA  4k3bA  4kwuA  4m00A  1bp1A  1cjsA  1ck1A  1ecrA  1f5qD  1fa9A  1gu7A  1itwA  1jkiA  1n80A  1nzjA  1qhdA  1qz9A  1sb7B  1vk1A  1vrmA  1xvuA  1yy3A  1z87A  2a1sC  2a3lA  2bt1A  2bydA  2c43A  2dfyC  2dlaA  2g3pA  2gg6A  2gsyE  2gzoA  2j2cA  2kfwA  2l9yA  2ntyB  2r3vA  2r58A  2w4mA  2x0cA  2y51A  2yb0E  2z86C  3afoA  3bu2A  3cvzA  3dupA  3eswA  3eukH  3fi7A  3fvvA  3gmsA  3hzzB  3m1uA  3mw8A  3mwcA  3nsjA  3ntkA  3oaaG  3ptyA  3rfyA  3seoB  3spgA  3u0kA  3vlaA  3vstA  4aqfB  4b21A  4dt4A  4dtfA  4ewtA  4f23A  4fzbC  4g1pA  4gfqA  4hvzA  4il6B  4jxkA  4m8mB  4mzyA  4onyA  4pyhA  4rg1A | 0.92  0.61  0.62  0.53  0.90  0.65  0.84  0.42  0.42  0.73  0.62  0.42  0.37  0.46  0.39  0.44  0.41  0.41  0.89  0.44  0.40  0.63  0.33  0.48  0.57  0.74  0.75  0.38  0.34  0.39  0.50  0.49  0.61  0.58  0.49  0.67  0.41  0.48  0.82  0.91  0.40  0.80  0.92  0.79  0.48  0.63  0.37  0.88  0.43  0.85  0.50  0.48  0.65  0.86  0.52  0.40  0.41  0.44  0.50  0.51  0.51  0.50  0.51  0.41  0.90  0.56  0.49  0.47  0.44  0.43  0.42  0.70  0.92  0.74  0.47  0.86  0.59  0.37  0.70  0.50  0.43  0.49  0.53  0.59  0.63  0.77  0.88  0.79  0.48  0.64  0.89  0.50  0.82  0.76  0.68  0.39  0.46  0.83  0.57  0.78  0.49  0.63  0.85  0.62  0.40  0.81  0.89  0.69  0.81  0.68  0.46  0.54  0.90  0.73  0.63  0.75  0.58  0.54 | 0.77  0.47  0.60  0.51  0.88  0.54  0.82  0.38  0.41  0.58  0.50  0.36  0.35  0.44  0.39  0.43  0.40  0.40  0.66  0.45  0.39  0.56  0.31  0.48  0.39  0.43  0.51  0.38  0.33  0.38  0.37  0.47  0.39  0.54  0.36  0.66  0.37  0.45  0.78  0.81  0.38  0.77  0.75  0.77  0.56  0.54  0.36  0.83  0.40  0.82  0.49  0.42  0.51  0.70  0.44  0.40  0.40  0.44  0.44  0.46  0.45  0.46  0.50  0.40  0.86  0.48  0.42  0.49  0.43  0.41  0.41  0.68  0.70  0.72  0.44  0.84  0.52  0.37  0.63  0.47  0.41  0.47  0.46  0.57  0.58  0.72  0.88  0.76  0.41  0.63  0.88  0.47  0.71  0.57  0.45  0.38  0.45  0.56  0.46  0.75  0.47  0.57  0.81  0.57  0.39  0.80  0.80  0.50  0.78  0.59  0.44  0.45  0.86  0.58  0.65  0.74  0.54  0.50 |

**Table S7.** Under the sequence identity cutoff of 70%, average coverage scores of 2dom proteins, 3dom proteins, m4dom proteins, 2dis proteins and all test protein in MPDB and DEMO-lib. *P*-value is determined by Student’s t-tests.

| Method | 2dom (166) | 3dom (69) | m4dom (40) | 2dis (81) | Total (356) | *P*-value |
| --- | --- | --- | --- | --- | --- | --- |
| MPDB | 0.61 | 0.61 | 0.54 | 0.62 | 0.61 | NA |
| DEMO-lib | 0.56 | 0.55 | 0.47 | 0.57 | 0.55 | 4.87E-06 |

**Table S8.** TM-score between the structural analogue with the highest *G*_score_ and the native structure of test proteins under the sequence identity cutoff of 30%, 50% and 70%, respectively.

| PDB ID | TM-score | | | PDB ID | TM-score | | | PDB ID | TM-score | | |
| --- | --- | --- | --- | --- | --- | --- | --- | --- | --- | --- | --- |
|  | 30% | 50% | 70% |  | 30% | 50% | 70% |  | 30% | 50% | 70% |
| 1cjyA  1efdN  1fjrA  1g87B  1hx6B  1iwaA  1m5qH  1mkfA  1mkmB  1nh2D  1pprM  1prrA  1q19A  1qwrA  1r71B  1rh1A  1rktA  1s6lA  1sp3A  1vz6A  1w3aA  1wv3A  1x7pA  1x9yA  1y11A  1yiqA  1zbuB  1ze1A  2ablA  2ahvA  2bkpA  2c1yA  2cxcA  2d1cA  2d7iA  2e9hA  2e9xB  2evrA  2ew9A  2fd5A  2gh8A  2gt1A  2gzaC  2hjqA  2hwjA  2ijd1  2iu7A  2iw2A  2jz4A  2kdyA  2kn4A  2mbgA  2nsfA  2nykA  2o6yA  2owbA  2qfiA  2qp2A  2qygA  2r5wB  2uu7A  2w4bA  2x7iA  2x8kC  2yilA  2yrqA  2zxcA  3a1iA  3a45A  3a56A  3ajvA  3aqkA  3arbA  3aujG  3b2zF  3b7wA  3bt3A  3c4tA  3craA  3d30A  3eo5A  3errA  3g79A  3h2tA  3hcsA  3hyiA  3i2dA  3iam2  3ifrA  3isqA  3j7aK  3k1rA  3k2iA  3kh5A  3kjpA  3kt1A  3ktmE  3kzwA  3l76A  3ld1A  3lsgA  3me4A  3ml4C  3mx2B  3mzfA  3njaB  3nqiA  3nt8A  3og5A  3oh0A  3pcsB  3po3S  3pxpA  3qavA  3qf4B  3qjjA  3qtdA  3r6bA  3rh7A | 0.25  0.90  0.33  0.61  0.70  0.87  0.39  0.59  0.64  0.47  0.39  0.43  0.84  0.84  0.78  0.45  0.71  0.32  0.35  0.33  0.39  0.76  0.70  0.37  0.49  0.45  0.46  0.81  0.48  0.77  0.44  0.58  0.74  0.59  0.35  0.45  0.73  0.76  0.54  0.82  0.59  0.84  0.48  0.33  0.45  0.67  0.49  0.82  0.30  0.52  0.47  0.61  0.31  0.71  0.87  0.91  0.36  0.50  0.90  0.30  0.79  0.30  0.82  0.82  0.46  0.37  0.26  0.75  0.80  0.43  0.73  0.78  0.86  0.35  0.64  0.78  0.74  0.59  0.35  0.81  0.27  0.70  0.80  0.35  0.45  0.51  0.73  0.74  0.92  0.86  0.86  0.41  0.49  0.84  0.57  0.70  0.32  0.89  0.58  0.34  0.80  0.60  0.47  0.29  0.79  0.45  0.68  0.34  0.80  0.58  0.66  0.37  0.29  0.81  0.86  0.85  0.94  0.32  0.45 | 0.81  0.91  0.36  0.94  0.70  0.93  0.39  0.59  0.64  0.95  0.90  0.43  0.84  0.84  0.78  0.45  0.71  0.38  0.35  0.33  0.39  0.76  0.79  0.37  0.49  0.92  0.55  0.78  0.58  0.77  0.41  0.57  0.74  0.59  0.74  0.42  0.73  0.81  0.54  0.82  0.77  0.84  0.51  0.38  0.37  0.70  0.84  0.82  0.47  0.52  0.35  0.61  0.31  0.71  0.95  0.92  0.66  0.50  0.90  0.91  0.88  0.87  0.82  0.82  0.57  0.37  0.93  0.75  0.80  0.43  0.78  0.78  0.93  0.42  0.85  0.78  0.74  0.59  0.35  0.84  0.36  0.70  0.75  0.34  0.36  0.51  0.77  0.89  0.93  0.88  0.96  0.41  0.49  0.84  0.57  0.70  0.32  0.90  0.63  0.34  0.94  0.65  0.47  0.32  0.82  0.45  0.68  0.34  0.81  0.58  0.66  0.72  0.29  0.81  0.86  0.88  0.94  0.78  0.45 | 0.81  0.91  0.36  0.94  0.70  0.99  0.39  0.59  0.64  0.95  0.90  0.43  0.84  0.84  0.78  0.45  0.71  0.38  0.35  0.52  0.39  0.76  0.79  0.37  0.49  0.94  0.55  0.78  0.58  0.77  0.41  0.57  0.74  0.59  0.74  0.42  0.73  0.81  0.54  0.82  0.94  0.84  0.51  0.38  0.37  0.70  0.98  0.82  0.47  0.64  0.46  0.61  0.31  0.71  0.95  0.95  0.66  0.50  0.99  0.91  0.92  0.87  0.82  0.82  0.57  0.37  0.93  0.75  0.80  0.43  0.78  0.78  0.97  0.95  0.92  0.78  0.74  0.59  0.35  0.84  0.36  0.70  0.75  0.34  0.88  0.51  0.77  0.89  0.93  0.88  0.98  0.41  0.95  0.84  0.57  0.70  0.32  0.90  0.63  0.34  0.94  0.65  0.47  0.32  0.92  0.45  0.68  0.34  0.81  0.58  0.66  0.72  0.29  0.81  0.86  0.88  0.94  0.78  0.45 | 3rwxA  3sb4A  3swjA  3t58B  3t7jA  3u07C  3u0oB  3u9gA  3ub1D  3uitD  3uo3A  3v7oB  3vr8B  3wkuA  3zvmA  4acoA  4ap5A  4axdA  4bfiB  4bt9B  4cczA  4d0nB  4d1iG  4dj3A  4dqaA  4eo3A  4eogA  4etxA  4fguA  4fkcA  4fxkC  4gbyA  4ggmX  4gslA  4gyjA  4h3tA  4hmoA  4ie6A  4l5gA  4lpqA  4m8rA  4n06B  4nj5A  4opaB  4qkuB  4up9A  4w7sA  1bf2A  1bhgA  1f7uA  1fx7A  1griA  1h88C  1m8pB  1ni5A  1q25A  1uzjA  1zpuA  1zy9A  2b5uA  2ewfA  2piaA  2r7dA  2uwnA  2v0nA  2vgmA  2wqrB  2y25B  2yk0A  2zzqA  3bt1U  3c1yA  3cw2C  3f83A  3fc3A  3gbgA  3h5cB  3ibjA  3ippB  3jymB  3kbgA  3mc8A  3npfA  3orjA  3plaA  3qe9Y  3qjoA  3qphA  3qyeA  3rimA  3rrpA  3soaA  3tixD  3tp9A  3ua3A  3uj0A  3vn4A  3vsmA  3w1bA  3zh9B  4alzA  4ax8A  4b3iA  4bd9B  4c0aB  4c0sA  4dimA  4indA  4jdzB  4kc3B  4kikB  4lmfA  4lziA  4m9pA  4pt5A  4uwhA  1c1zA  1d2pA  1k7tA | 0.31  0.45  0.53  0.48  0.69  0.67  0.80  0.33  0.34  0.31  0.76  0.40  0.87  0.29  0.32  0.72  0.69  0.32  0.78  0.40  0.75  0.77  0.69  0.79  0.35  0.43  0.77  0.33  0.39  0.88  0.83  0.87  0.38  0.37  0.82  0.74  0.76  0.35  0.57  0.49  0.48  0.86  0.25  0.27  0.67  0.73  0.52  0.76  0.84  0.85  0.75  0.42  0.35  0.62  0.61  0.44  0.38  0.91  0.77  0.47  0.35  0.61  0.86  0.45  0.27  0.51  0.58  0.49  0.45  0.41  0.35  0.44  0.66  0.39  0.27  0.36  0.79  0.47  0.39  0.51  0.25  0.49  0.75  0.57  0.58  0.75  0.66  0.41  0.75  0.83  0.77  0.56  0.32  0.38  0.41  0.71  0.36  0.76  0.64  0.58  0.60  0.36  0.84  0.32  0.46  0.49  0.78  0.40  0.59  0.72  0.65  0.46  0.27  0.32  0.77  0.86  0.49  0.38  0.39 | 0.31  0.45  0.53  0.48  0.69  0.67  0.89  0.27  0.34  0.48  0.76  0.40  0.89  0.29  0.32  0.72  0.93  0.77  0.78  0.40  0.75  0.82  0.69  0.79  0.35  0.43  0.77  0.33  0.91  0.89  0.83  0.87  0.37  0.37  0.82  0.90  0.76  0.35  0.57  0.49  0.48  0.86  0.86  0.38  0.67  0.88  0.52  0.85  0.92  0.84  0.75  0.42  0.64  0.62  0.61  0.44  0.63  0.90  0.77  0.47  0.35  0.70  0.86  0.45  0.35  0.57  0.53  0.49  0.37  0.41  0.35  0.44  0.65  0.55  0.35  0.36  0.79  0.47  0.92  0.51  0.90  0.49  0.75  0.57  0.60  0.75  0.84  0.41  0.82  0.94  0.93  0.56  0.44  0.38  0.48  0.74  0.36  0.76  0.64  0.58  0.60  0.36  0.88  0.32  0.59  0.81  0.78  0.40  0.59  0.72  0.65  0.65  0.27  0.32  0.87  0.86  0.44  0.38  0.39 | 0.31  0.45  0.91  0.48  0.69  0.67  0.89  0.27  0.34  0.48  0.76  0.40  0.94  0.29  0.32  0.72  0.93  0.77  0.78  0.40  0.75  0.82  0.90  0.85  0.35  0.43  0.77  0.33  0.91  0.93  0.83  0.87  0.37  0.37  0.82  0.90  0.76  0.35  0.57  0.49  0.48  0.86  0.86  0.38  0.94  0.90  0.52  0.85  0.92  0.84  0.93  0.42  0.64  0.62  0.61  0.44  0.63  0.90  0.77  0.47  0.35  0.70  0.86  0.45  0.35  0.57  0.53  0.49  0.37  0.41  0.86  0.44  0.65  0.55  0.35  0.36  0.79  0.47  0.92  0.51  0.90  0.74  0.75  0.57  0.60  0.75  0.84  0.67  0.92  0.94  0.93  0.56  0.44  0.38  0.48  0.74  0.36  0.76  0.64  0.58  0.60  0.36  0.88  0.32  0.59  0.53  0.78  0.40  0.59  0.72  0.86  0.65  0.27  0.32  0.87  0.90  0.44  0.38  0.39 | 1kfqA  1ldjA  1nyqB  1ug9A  1z1wA  2au3A  2ii2A  2olsA  2ra1A  2v5dA  2xt6A  2zpaB  3apoA  3b43A  3gf5B  3hjlA  3kq4B  3kwlA  3ob8A  3opfB  3p53A  3pvlA  3r05A  3ubhA  3w2wA  3zniA  4aimA  4ak1A  4aq1A  4fe9A  4h2aA  4i5sB  4iggB  4j9vA  4k3bA  4kwuA  4m00A  1bp1A  1cjsA  1ck1A  1ecrA  1f5qD  1fa9A  1gu7A  1itwA  1jkiA  1n80A  1nzjA  1qhdA  1qz9A  1sb7B  1vk1A  1vrmA  1xvuA  1yy3A  1z87A  2a1sC  2a3lA  2bt1A  2bydA  2c43A  2dfyC  2dlaA  2g3pA  2gg6A  2gsyE  2gzoA  2j2cA  2kfwA  2l9yA  2ntyB  2r3vA  2r58A  2w4mA  2x0cA  2y51A  2yb0E  2z86C  3afoA  3bu2A  3cvzA  3dupA  3eswA  3eukH  3fi7A  3fvvA  3gmsA  3hzzB  3m1uA  3mw8A  3mwcA  3nsjA  3ntkA  3oaaG  3ptyA  3rfyA  3seoB  3spgA  3u0kA  3vlaA  3vstA  4aqfB  4b21A  4dt4A  4dtfA  4ewtA  4f23A  4fzbC  4g1pA  4gfqA  4hvzA  4il6B  4jxkA  4m8mB  4mzyA  4onyA  4pyhA  4rg1A | 0.77  0.42  0.29  0.52  0.87  0.65  0.81  0.41  0.26  0.58  0.26  0.82  0.28  0.35  0.21  0.33  0.36  0.41  0.52  0.26  0.43  0.40  0.29  0.46  0.62  0.27  0.31  0.18  0.25  0.30  0.29  0.44  0.62  0.61  0.38  0.63  0.31  0.72  0.83  0.83  0.33  0.83  0.52  0.79  0.48  0.70  0.30  0.75  0.52  0.92  0.78  0.73  0.86  0.86  0.30  0.30  0.37  0.51  0.70  0.72  0.74  0.46  0.62  0.31  0.88  0.31  0.61  0.43  0.43  0.36  0.38  0.69  0.25  0.71  0.59  0.85  0.36  0.33  0.75  0.41  0.32  0.48  0.42  0.61  0.66  0.81  0.90  0.74  0.54  0.63  0.89  0.37  0.80  0.80  0.36  0.34  0.32  0.78  0.51  0.68  0.56  0.34  0.83  0.75  0.32  0.76  0.59  0.82  0.81  0.63  0.28  0.48  0.84  0.23  0.73  0.64  0.53  0.34 | 0.89  0.87  0.87  0.52  0.87  0.65  0.81  0.41  0.33  0.72  0.75  0.82  0.28  0.35  0.30  0.33  0.36  0.35  0.89  0.55  0.43  0.84  0.29  0.47  0.62  0.71  0.31  0.18  0.25  0.33  0.70  0.44  0.62  0.61  0.54  0.63  0.31  0.74  0.83  0.89  0.43  0.83  0.95  0.86  0.48  0.94  0.30  0.89  0.52  0.92  0.78  0.73  0.91  0.86  0.78  0.30  0.39  0.51  0.70  0.72  0.74  0.52  0.62  0.34  0.95  0.90  0.65  0.82  0.43  0.36  0.38  0.69  0.64  0.71  0.59  0.94  0.81  0.33  0.74  0.50  0.32  0.49  0.77  0.61  0.77  0.81  0.90  0.84  0.90  0.67  0.89  0.37  0.80  0.85  0.91  0.35  0.32  0.80  0.52  0.87  0.56  0.74  0.90  0.75  0.32  0.81  0.90  0.82  0.94  0.65  0.28  0.48  0.93  0.80  0.83  0.93  0.53  0.80 | 0.91  0.87  0.87  0.52  0.87  0.65  0.81  0.41  0.33  0.72  0.75  0.82  0.28  0.35  0.30  0.33  0.36  0.35  0.89  0.55  0.43  0.84  0.29  0.47  0.62  0.71  0.77  0.18  0.25  0.33  0.70  0.44  0.62  0.61  0.54  0.63  0.31  0.74  0.83  0.92  0.43  0.83  0.96  0.86  0.48  0.94  0.30  0.89  0.52  0.92  0.78  0.73  0.91  0.86  0.78  0.30  0.39  0.51  0.70  0.72  0.74  0.52  0.62  0.34  0.95  0.90  0.65  0.82  0.43  0.36  0.38  0.69  0.98  0.71  0.59  0.94  0.81  0.33  0.74  0.50  0.32  0.49  0.77  0.61  0.77  0.81  0.90  0.93  0.90  0.67  0.89  0.37  0.80  0.85  0.91  0.35  0.32  0.96  0.55  0.87  0.56  0.90  0.90  0.77  0.32  0.81  0.90  0.82  0.97  0.65  0.28  0.48  0.93  0.80  0.83  0.93  0.53  0.80 |

**Table S9.** Under the sequence identity (SeqId) cutoff of 30%, 50% and 70%, average TM-score of the structural analogue with the highest *G*_score_. #TM-score > 0.5 represents the number of the structural analogues with TM-score > 0.5.

| SeqId cutoff | 2dom (166) | 3dom (69) | m4dom (40) | 2dis (81) | Total (356) | #TM-score > 0.5 |
| --- | --- | --- | --- | --- | --- | --- |
| 30% | 0.58 | 0.55 | 0.44 | 0.59 | 0.56 | 192 |
| 50% | 0.65 | 0.61 | 0.53 | 0.69 | 0.64 | 239 |
| 70% | 0.67 | 0.63 | 0.55 | 0.70 | 0.65 | 247 |

**Table S10.** The *G*_score_ of the top 1 structural analogue detected under the sequence identity cutoff of 30%, 50% and 70%, respectively.

| PDB ID | *G*_score_ | | | PDB ID | *G*_score_ | | | PDB ID | *G*_score_ | | |
| --- | --- | --- | --- | --- | --- | --- | --- | --- | --- | --- | --- |
|  | 30% | 50% | 70% |  | 30% | 50% | 70% |  | 30% | 50% | 70% |
| 1cjyA  1efdN  1fjrA  1g87B  1hx6B  1iwaA  1m5qH  1mkfA  1mkmB  1nh2D  1pprM  1prrA  1q19A  1qwrA  1r71B  1rh1A  1rktA  1s6lA  1sp3A  1vz6A  1w3aA  1wv3A  1x7pA  1x9yA  1y11A  1yiqA  1zbuB  1ze1A  2ablA  2ahvA  2bkpA  2c1yA  2cxcA  2d1cA  2d7iA  2e9hA  2e9xB  2evrA  2ew9A  2fd5A  2gh8A  2gt1A  2gzaC  2hjqA  2hwjA  2ijd1  2iu7A  2iw2A  2jz4A  2kdyA  2kn4A  2mbgA  2nsfA  2nykA  2o6yA  2owbA  2qfiA  2qp2A  2qygA  2r5wB  2uu7A  2w4bA  2x7iA  2x8kC  2yilA  2yrqA  2zxcA  3a1iA  3a45A  3a56A  3ajvA  3aqkA  3arbA  3aujG  3b2zF  3b7wA  3bt3A  3c4tA  3craA  3d30A  3eo5A  3errA  3g79A  3h2tA  3hcsA  3hyiA  3i2dA  3iam2  3ifrA  3isqA  3j7aK  3k1rA  3k2iA  3kh5A  3kjpA  3kt1A  3ktmE  3kzwA  3l76A  3ld1A  3lsgA  3me4A  3ml4C  3mx2B  3mzfA  3njaB  3nqiA  3nt8A  3og5A  3oh0A  3pcsB  3po3S  3pxpA  3qavA  3qf4B  3qjjA  3qtdA  3r6bA  3rh7A | 0.46  0.86  0.41  0.60  0.69  0.82  0.59  0.53  0.84  0.65  0.46  0.73  0.76  0.79  0.71  0.40  0.74  0.49  0.39  0.43  0.55  0.78  0.78  0.47  0.52  0.58  0.52  0.80  0.75  0.77  0.57  0.55  0.75  0.48  0.46  0.48  0.71  0.71  0.82  0.78  0.72  0.80  0.72  0.55  0.49  0.48  0.50  0.79  0.50  0.51  0.57  0.53  0.47  0.72  0.88  0.91  0.48  0.58  0.82  0.50  0.72  0.42  0.76  0.77  0.71  0.64  0.38  0.58  0.78  0.46  0.67  0.73  0.88  0.46  0.78  0.91  0.70  0.77  0.55  0.80  0.45  0.58  0.83  0.45  0.45  0.52  0.79  0.73  0.90  0.83  0.81  0.55  0.55  0.81  0.67  0.69  0.50  0.83  0.58  0.41  0.84  0.75  0.78  0.39  0.82  0.74  0.68  0.41  0.82  0.59  0.66  0.51  0.47  0.83  0.87  0.82  0.92  0.51  0.47 | 0.87  0.87  0.43  0.93  0.69  0.92  0.59  0.53  0.84  0.92  0.87  0.73  0.76  0.79  0.71  0.40  0.74  0.49  0.39  0.43  0.55  0.78  0.84  0.47  0.52  0.83  0.54  0.81  0.83  0.77  0.59  0.83  0.75  0.48  0.84  0.51  0.71  0.75  0.82  0.78  0.83  0.80  0.82  0.55  0.49  0.49  0.81  0.79  0.74  0.51  0.57  0.53  0.47  0.72  0.92  0.92  0.71  0.58  0.82  0.90  0.81  0.86  0.77  0.77  0.72  0.64  0.91  0.59  0.78  0.46  0.70  0.73  0.93  0.47  0.86  0.91  0.70  0.77  0.55  0.84  0.51  0.58  0.85  0.46  0.50  0.52  0.83  0.85  0.91  0.86  0.93  0.55  0.55  0.81  0.67  0.69  0.50  0.84  0.60  0.41  0.90  0.79  0.78  0.39  0.88  0.74  0.68  0.41  0.88  0.59  0.66  0.65  0.47  0.83  0.89  0.86  0.92  0.82  0.47 | 0.87  0.87  0.43  0.93  0.69  0.99  0.59  0.53  0.84  0.92  0.87  0.73  0.76  0.79  0.71  0.40  0.74  0.49  0.39  0.48  0.55  0.78  0.84  0.47  0.52  0.90  0.54  0.81  0.83  0.77  0.59  0.83  0.75  0.48  0.84  0.51  0.71  0.75  0.82  0.78  0.94  0.80  0.82  0.55  0.49  0.49  0.98  0.79  0.74  0.71  0.73  0.53  0.47  0.72  0.92  0.93  0.71  0.58  0.97  0.90  0.89  0.86  0.77  0.77  0.72  0.64  0.91  0.59  0.78  0.46  0.70  0.73  0.96  0.89  0.89  0.91  0.70  0.77  0.55  0.84  0.51  0.58  0.85  0.46  0.80  0.52  0.83  0.85  0.91  0.86  0.95  0.55  0.95  0.81  0.67  0.69  0.50  0.84  0.60  0.41  0.90  0.79  0.78  0.39  0.95  0.74  0.68  0.41  0.88  0.59  0.66  0.65  0.47  0.83  0.89  0.86  0.92  0.82  0.47 | 3rwxA  3sb4A  3swjA  3t58B  3t7jA  3u07C  3u0oB  3u9gA  3ub1D  3uitD  3uo3A  3v7oB  3vr8B  3wkuA  3zvmA  4acoA  4ap5A  4axdA  4bfiB  4bt9B  4cczA  4d0nB  4d1iG  4dj3A  4dqaA  4eo3A  4eogA  4etxA  4fguA  4fkcA  4fxkC  4gbyA  4ggmX  4gslA  4gyjA  4h3tA  4hmoA  4ie6A  4l5gA  4lpqA  4m8rA  4n06B  4nj5A  4opaB  4qkuB  4up9A  4w7sA  1bf2A  1bhgA  1f7uA  1fx7A  1griA  1h88C  1m8pB  1ni5A  1q25A  1uzjA  1zpuA  1zy9A  2b5uA  2ewfA  2piaA  2r7dA  2uwnA  2v0nA  2vgmA  2wqrB  2y25B  2yk0A  2zzqA  3bt1U  3c1yA  3cw2C  3f83A  3fc3A  3gbgA  3h5cB  3ibjA  3ippB  3jymB  3kbgA  3mc8A  3npfA  3orjA  3plaA  3qe9Y  3qjoA  3qphA  3qyeA  3rimA  3rrpA  3soaA  3tixD  3tp9A  3ua3A  3uj0A  3vn4A  3vsmA  3w1bA  3zh9B  4alzA  4ax8A  4b3iA  4bd9B  4c0aB  4c0sA  4dimA  4indA  4jdzB  4kc3B  4kikB  4lmfA  4lziA  4m9pA  4pt5A  4uwhA  1c1zA  1d2pA  1k7tA | 0.61  0.67  0.79  0.60  0.64  0.79  0.79  0.50  0.60  0.52  0.75  0.58  0.83  0.43  0.57  0.63  0.66  0.40  0.81  0.61  0.80  0.86  0.54  0.74  0.64  0.48  0.72  0.64  0.47  0.85  0.80  0.84  0.56  0.42  0.77  0.73  0.77  0.50  0.80  0.77  0.65  0.82  0.40  0.50  0.54  0.70  0.74  0.65  0.77  0.77  0.74  0.76  0.56  0.88  0.76  0.80  0.45  0.86  0.69  0.43  0.50  0.75  0.70  0.64  0.50  0.76  0.82  0.71  0.49  0.51  0.42  0.47  0.82  0.55  0.47  0.49  0.73  0.75  0.63  0.65  0.51  0.72  0.73  0.73  0.49  0.49  0.46  0.48  0.66  0.80  0.69  0.54  0.43  0.48  0.46  0.71  0.42  0.69  0.81  0.75  0.69  0.48  0.84  0.47  0.51  0.77  0.79  0.62  0.78  0.83  0.79  0.54  0.55  0.75  0.82  0.82  0.66  0.61  0.36 | 0.61  0.67  0.79  0.60  0.64  0.79  0.86  0.53  0.60  0.74  0.75  0.58  0.87  0.43  0.57  0.63  0.92  0.71  0.81  0.61  0.80  0.86  0.54  0.74  0.64  0.48  0.72  0.64  0.86  0.88  0.81  0.84  0.57  0.42  0.77  0.82  0.77  0.50  0.80  0.77  0.65  0.82  0.84  0.51  0.54  0.89  0.82  0.77  0.87  0.79  0.74  0.76  0.72  0.88  0.76  0.80  0.67  0.87  0.69  0.43  0.50  0.90  0.70  0.64  0.52  0.76  0.82  0.71  0.53  0.51  0.42  0.47  0.83  0.69  0.47  0.49  0.73  0.75  0.87  0.65  0.85  0.72  0.73  0.73  0.88  0.49  0.63  0.48  0.67  0.92  0.88  0.54  0.44  0.48  0.82  0.72  0.42  0.69  0.81  0.75  0.69  0.48  0.88  0.53  0.74  0.85  0.79  0.62  0.78  0.83  0.79  0.88  0.55  0.75  0.87  0.83  0.69  0.61  0.41 | 0.61  0.67  0.85  0.60  0.64  0.79  0.86  0.53  0.60  0.74  0.75  0.58  0.94  0.43  0.57  0.63  0.92  0.71  0.81  0.61  0.80  0.86  0.83  0.85  0.64  0.48  0.72  0.64  0.86  0.90  0.81  0.84  0.57  0.42  0.77  0.82  0.77  0.50  0.80  0.77  0.65  0.82  0.84  0.51  0.88  0.90  0.82  0.77  0.87  0.79  0.91  0.76  0.72  0.88  0.76  0.80  0.67  0.87  0.69  0.43  0.50  0.90  0.70  0.64  0.52  0.76  0.82  0.71  0.53  0.51  0.84  0.47  0.83  0.69  0.47  0.49  0.73  0.75  0.87  0.65  0.85  0.84  0.73  0.73  0.88  0.49  0.63  0.49  0.90  0.92  0.88  0.54  0.44  0.48  0.82  0.72  0.42  0.69  0.81  0.75  0.69  0.48  0.88  0.53  0.74  0.88  0.79  0.62  0.78  0.83  0.90  0.88  0.55  0.75  0.87  0.90  0.69  0.61  0.41 | 1kfqA  1ldjA  1nyqB  1ug9A  1z1wA  2au3A  2ii2A  2olsA  2ra1A  2v5dA  2xt6A  2zpaB  3apoA  3b43A  3gf5B  3hjlA  3kq4B  3kwlA  3ob8A  3opfB  3p53A  3pvlA  3r05A  3ubhA  3w2wA  3zniA  4aimA  4ak1A  4aq1A  4fe9A  4h2aA  4i5sB  4iggB  4j9vA  4k3bA  4kwuA  4m00A  1bp1A  1cjsA  1ck1A  1ecrA  1f5qD  1fa9A  1gu7A  1itwA  1jkiA  1n80A  1nzjA  1qhdA  1qz9A  1sb7B  1vk1A  1vrmA  1xvuA  1yy3A  1z87A  2a1sC  2a3lA  2bt1A  2bydA  2c43A  2dfyC  2dlaA  2g3pA  2gg6A  2gsyE  2gzoA  2j2cA  2kfwA  2l9yA  2ntyB  2r3vA  2r58A  2w4mA  2x0cA  2y51A  2yb0E  2z86C  3afoA  3bu2A  3cvzA  3dupA  3eswA  3eukH  3fi7A  3fvvA  3gmsA  3hzzB  3m1uA  3mw8A  3mwcA  3nsjA  3ntkA  3oaaG  3ptyA  3rfyA  3seoB  3spgA  3u0kA  3vlaA  3vstA  4aqfB  4b21A  4dt4A  4dtfA  4ewtA  4f23A  4fzbC  4g1pA  4gfqA  4hvzA  4il6B  4jxkA  4m8mB  4mzyA  4onyA  4pyhA  4rg1A | 0.68  0.60  0.49  0.45  0.88  0.55  0.84  0.73  0.54  0.62  0.45  0.88  0.40  0.82  0.41  0.50  0.61  0.45  0.49  0.40  0.84  0.45  0.36  0.80  0.52  0.47  0.42  0.55  0.54  0.44  0.41  0.55  0.72  0.72  0.49  0.47  0.54  0.66  0.98  0.80  0.50  0.80  0.52  0.77  0.47  0.73  0.39  0.71  0.51  0.90  0.83  0.73  0.84  0.76  0.48  0.54  0.49  0.45  0.69  0.70  0.71  0.65  0.64  0.40  0.85  0.48  0.56  0.43  0.59  0.50  0.49  0.73  0.48  0.73  0.86  0.82  0.48  0.48  0.72  0.52  0.50  0.49  0.47  0.55  0.67  0.75  0.88  0.73  0.52  0.80  0.86  0.45  0.78  0.76  0.48  0.46  0.49  0.75  0.55  0.66  0.54  0.42  0.80  0.78  0.44  0.83  0.51  0.79  0.84  0.79  0.51  0.44  0.83  0.39  0.73  0.75  0.70  0.44 | 0.90  0.82  0.90  0.45  0.88  0.55  0.84  0.73  0.58  0.75  0.55  0.88  0.40  0.82  0.44  0.50  0.65  0.44  0.82  0.83  0.84  0.88  0.36  0.82  0.52  0.48  0.42  0.55  0.54  0.46  0.71  0.55  0.72  0.72  0.86  0.47  0.54  0.84  0.98  0.87  0.50  0.80  0.94  0.86  0.47  0.93  0.39  0.85  0.51  0.90  0.83  0.73  0.87  0.76  0.77  0.54  0.50  0.45  0.69  0.70  0.71  0.81  0.64  0.42  0.94  0.86  0.60  0.81  0.60  0.50  0.49  0.73  0.85  0.73  0.86  0.92  0.79  0.48  0.74  0.57  0.50  0.50  0.75  0.55  0.82  0.75  0.88  0.84  0.87  0.88  0.87  0.45  0.78  0.82  0.88  0.47  0.49  0.80  0.84  0.83  0.54  0.81  0.87  0.78  0.44  0.92  0.90  0.79  0.94  0.92  0.52  0.44  0.93  0.80  0.83  0.94  0.70  0.75 | 0.91  0.82  0.90  0.45  0.88  0.55  0.84  0.73  0.58  0.75  0.55  0.88  0.40  0.82  0.44  0.50  0.65  0.44  0.82  0.83  0.84  0.88  0.36  0.82  0.52  0.48  0.81  0.55  0.54  0.46  0.71  0.55  0.72  0.72  0.86  0.47  0.54  0.84  0.98  0.90  0.50  0.80  0.95  0.86  0.47  0.93  0.39  0.85  0.51  0.90  0.83  0.73  0.87  0.76  0.77  0.54  0.50  0.45  0.69  0.70  0.71  0.81  0.64  0.42  0.94  0.86  0.60  0.81  0.60  0.50  0.49  0.73  0.98  0.73  0.86  0.92  0.79  0.48  0.74  0.57  0.50  0.50  0.75  0.55  0.82  0.75  0.88  0.93  0.87  0.88  0.87  0.45  0.78  0.82  0.88  0.47  0.49  0.94  0.90  0.83  0.54  0.91  0.87  0.78  0.44  0.92  0.90  0.79  0.96  0.92  0.52  0.44  0.93  0.80  0.83  0.94  0.70  0.75 |

**Supplementary Texts**

**Text S1. Sequence identity calculation.**

PSI-BLAST is used to calculate sequence identities. PSI-BLAST is used locally to build a position specific similarity matrix (PSSM) or profile from homologous sequences in NCBI’s non-redundant protein sequence database with each unique PDB sequence as query. Three iterations are performed for each query, with an E-value cutoff of 0.0001 for inclusion in the profile. We control for drift in the PSSM by checking to see whether hits in previous rounds with E-values better than 0.0001 appear with E-values worse than 0.0001 in subsequent rounds. If so, we take the last profile not exhibiting drift. The resulting matrix is used to search the PDB for sequences related to each query with an E-value better than 1.0 and alignment length greater than 20, and the resulting sequence identities and alignment lengths from the PSI-BLAST output are stored.
